# The Intrinsically Disordered Region of Coronins Fine-tunes Oligomerization and Actin Polymerization

**DOI:** 10.1101/2022.01.19.477021

**Authors:** Xiao Han, Zixin Hu, Wahyu Surya, Qianqian Ma, Feng Zhou, Lars Nordenskiöld, Jaume Torres, Lanyuan Lu, Yansong Miao

## Abstract

Coronins are highly conserved actin-binding proteins (ABPs) in the eukaryotic kingdom for polymerizing actin cytoskeleton. The biochemical activity of coronins is primarily mediated by the structural N-terminal β-propeller and the C-terminal helical coiled-coil (CC) domains, but less is known about the function of a middle nonconserved region, the “unique region (UR)”. The coronin UR is an intrinsically disordered region (IDR). Herein, we demonstrate that the low complexity of the UR is a conserved signature of the coronin protein family, and the UR/IDR exhibits a striking evolutionary correlated pattern associated with sequence length. By analyzing the role of the IDR in coronins via coarse-grained simulations, we reveal that evolutionary selection of IDR length is coupled with the oligomerization of IDR-containing proteins (IDPs) to provide optimal functional output. By integrating biochemical and cell biology experiments and protein engineering, we found that the IDR regulates Crn1 biochemical activity, both *in vivo* and *in vitro*, by fine-tuning CC domain oligomerization and maintaining Crn1 in a tetrameric state. The IDR-guided optimization of Crn1 oligomerization is critical for Arp2/3-mediated actin polymerization.

## Introduction

Coronins were first identified in *Dictyostelium*, in which they localize on “crown-shaped” F-actin structures at the dorsal cell surface ^1^. Subsequently, coronins were found to be highly conserved eukaryotic actin-binding proteins (ABPs) spanning from yeast to human that regulate actin cytoskeleton polymerization and remodeling ^2–6^. A lack of coronin results in defects in various cellular processes, such as cell motility ^7–15^, cytokinesis ^13,15^, endocytosis ^13,16–19^, immune cell function ^20–24^ and actin turnover ^7,13,25^. The inter- and intramolecular interactions of coronin proteins with F-actin and other ABPs are critical to regulate coronin function at different steps of actin polymerization, such as inhibition of actin nucleation through interaction with the Arp2/3 complex ^7–10,22,26–28^, actin bundling through direct association with F-actin ^1,10,29,30^ and actin depolymerization in cooperation with ADF/cofilin ^25,31–35^. Crn1 in budding yeast is functionally relevant to the Arp2/3 complex. Crn1 overexpression could rescue the growth defects of *arc35-26*, but a synthetic growth defect was observed when *CRN1* was deleted in the background of another Arp2/3 temperature-sensitive mutant, *arp2-21* ^26^. A previous structural study showed direct binding of Crn1 to the Arp2/3 complex near the p35/Arc35/ARPC2 subunit, which kept the Arp2/3 complex in an open (inactive) conformation ^27,28^. The mammalian coronin isoform Coronin 1A also functionally associates with the Arp2/3 complex, F-actin and cofilin ^25,36,37^. While Coronin 1A is more relevant to immune cells, mammalian Coronin 1B and Coronin 1C are ubiquitously expressed ^2^. Coronin1A was found to control actin equilibrium in T cells, in which its deletion induced an increase in F-actin but a decrease in the G-actin pool in an Arp2/3-dependent manner ^21^. Coronin 1C formed actin-dependent filamentous and punctate structures at the leading edge in human cells and was associated with F-actin *in vivo* ^38^. Similarly, Coronin 1B regulates cell motility due to its localization at the cell leading edge and lamellipodia and coordinately functions with the Arp2/3 complex in a coronin phosphorylation-dependent manner ^7,9,10^.

The biochemical activities of coronin family proteins *in vitro* and *in vivo* are known to be conferred by the family-conserved β-propeller and coiled-coil (CC) domains, located at the N- and C-terminus, respectively. Whereas the well-folded β-propeller domain primarily interacts with F-actin ^10,29,39^, the CC domains of different coronins carry out a diverse array of activities, including direct binding with F-actin, inhibition of the Arp2/3 complex and regulation of actin turnover, in cooperation with Cof1 ^26,29,31,36,38^. Although coronin CC domains play various roles in actin polymerization, they share a chemical-physical property, protein oligomerization ^36,38,40–43^. However, the specific oligomerization state of each coronin and how oligomerization is regulated are still unclear.

In budding yeast, both the N-terminal β-propeller (aa 1-400) and the CC domains (aa 600-651) of Crn1 are indispensable for maintaining functional localization on cortical endocytic actin patches ^3,26,44^. In the present work, we demonstrate that the Crn1 “unique region” (UR, aa 401-604), located between the β-propeller and CC domain, is a highly flexible intrinsically disordered region (IDR) that directly fine-tunes CC oligomerization and thereby coronin activities. Our evolutionary analysis shows that the presence of the UR (hereafter named the IDR) is a hallmark of coronin family members, although sequence and length are variable. The length of IDRs in coronin family proteins exhibits an intriguing evolutionary pattern; IDRs are long in coronins in the fungal kingdom and short in coronins in mammals. Based on a systematic biochemical and biophysical characterization of Crn1, we found that the IDR provides structural flexibility for neighboring β-propeller and CC domains to maintain proper protein conformation and *in vivo* localization. Strikingly, we further discovered that the IDR precisely controls Crn1 oligomerization and thereby function by tuning and optimizing the oligomeric state of the Crn1 CC domain. In statistical thermodynamics, the stable structure of a protein or protein complex is determined by the potential energy surface (i.e., the “energy landscape”) ^45,46^. Molecular dynamics (MD) simulations and analytical ultracentrifugation showed that the Crn1 IDR modulates the energy landscape of the Crn1 CC domain, inducing tetramer formation in an IDR length-dependent manner. Here, we refer to a one-dimensional energy landscape, taking the oligomeric number as the coordinate. We found that the length of the coronin IDR appears to have been evolutionarily selected depending on the packing pattern of the coronin CC domain involving hydrophobic residues. The IDR modulates the energetic stability of the CC domain during the formation of helical bundles. Once the mammalian CC domain has evolved an ideal heptad repeat in its sequence, perfect contact favored by stable hydrophobic interactions between adjacent helices in oligomeric CC domains will form, and the length of the IDR plays a less critical role in tuning the CC oligomeric state. Furthermore, we demonstrate the importance of appropriate oligomerization for coronin proteins, both *in vivo* and *in vitro*, via *de novo* design and protein engineering. We created Crn1 variants in different homo-oligomeric states. Quantitative biochemical analysis of actin polymerization and cell biology-based functionalization assays showed that tetrameric coronin is more functional than dimeric Crn1 variants with respect to actin bundling and Arp2/3 inhibition *in vitro* and genetic interaction with the Arp2/3 complex *in vivo*. Our work reveals that variations in IDR length during evolution can modulate the energy landscape of CC domain oligomerization. This is a novel mechanism by which the coronin IDR optimizes coronin activity by fine-tuning CC domain-mediated oligomerization.

## Results

### IDR is indispensable for Crn1 localization at the actin patch and its interaction with actin filaments

To characterize the IDR of Crn1, we first analyzed the Crn1 sequence using IUPred2 ^47,48^ and defined the Crn1-IDR (aa 401-604) between the N-terminal β-propeller domain (Crn1-N, aa 1-400) and C-terminal CC domain (Crn1-CC, aa 605-651) (Figure 1A) ^29^. To dissect the functional cooperation of the three Crn1 domains *in vivo*, we generated genome-integrating plasmids to express truncated variants of Crn1 C-terminally tagged with a monomeric red fluorescent protein (mRuby2) in *Δcrn1* under control of the endogenous Crn1 promoter (Figure 1B). Full-length Crn1 (Crn1-FL) localized to cortical actin patches, but all six truncated Crn1 variants that lacked one or two domains displayed a diffuse pattern without clear punctate actin patch signals (Figure 1C). The loss of actin patch localization in Crn1 variants without the N or CC region (Figure 1B) is consistent with their previously reported individual and synergistic function in F-actin binding ^26,29,31,39,49^. However, the complete loss of the patch pattern of Crn1-ΔIDR was rather unexpected. To examine whether the mislocalization of Crn1-ΔIDR was due to a change in protein expression or degradation, we characterized the expression of truncated Crn1 variants *in vivo*, and Crn1-ΔIDR showed a protein expression level similar to that of Crn1-FL (Figure 1D).

**Figure 1.**
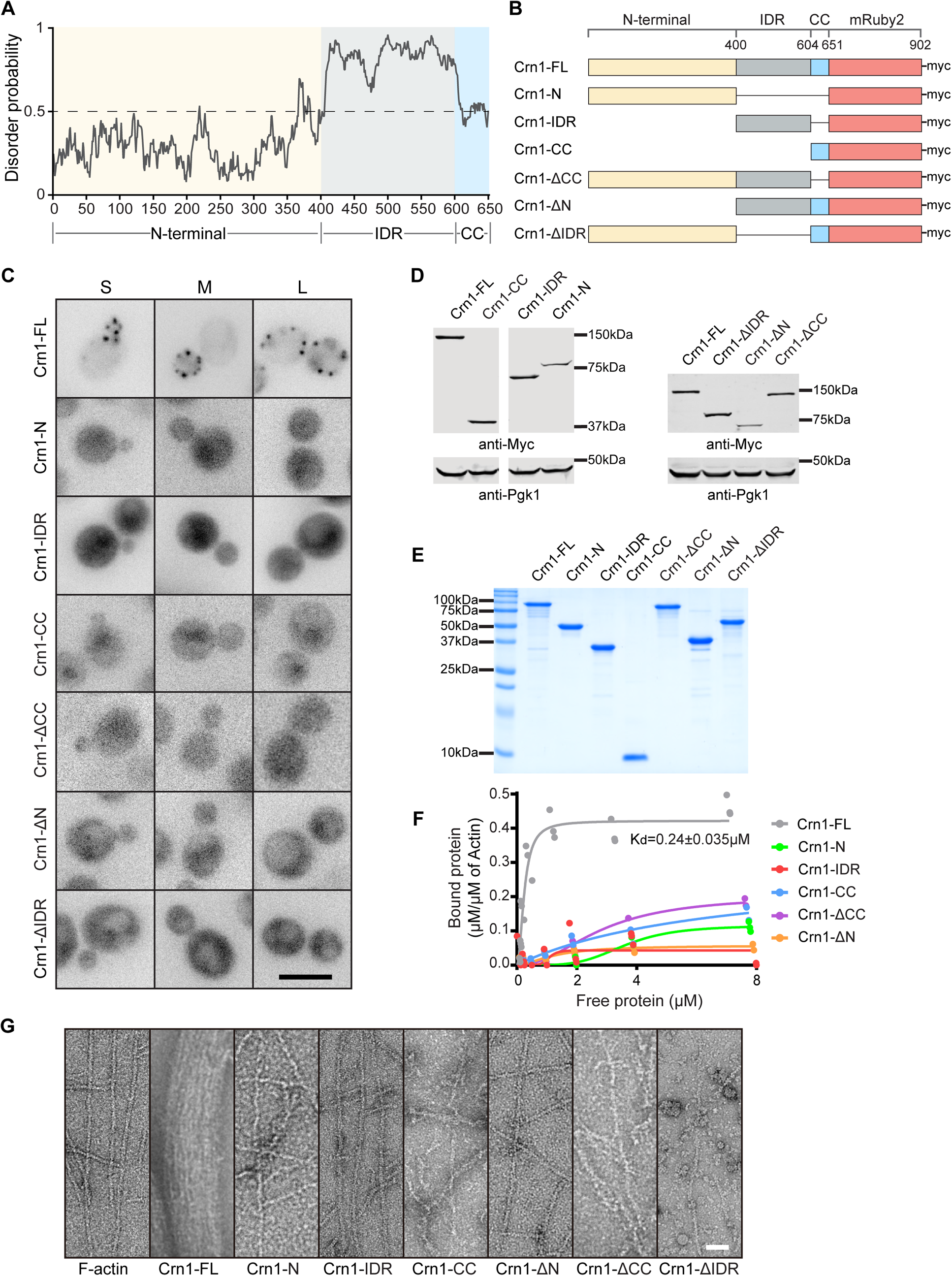
IDR is indispensable for Crn1 localization at the actin patch and its interaction with actin filaments. A. IDR prediction using IUPs (website: https://iupred2a.elte.hu/) for Crn1, with a disorder probability of 0.5 as the cutoff.
B. Schematic representation of yeast *in vivo* expression of full-length (FL) Crn1 and truncated Crn1 variants tagged with mRuby2 and 4xMyc tags at the C-terminus under the Crn1 native promoter.
C. Representative images of mRuby2-tagged Crn1-FL and truncated Crn1 variants *in vivo*. S, small-budded cell; M, middle-budded cell; L, large-budded cell. Scale bar, 5 µm.
D. Western blot analysis of yeast Crn1 expression in (C) using antibodies against the Myc tag. Anti-Pgk1 was used as a loading control.
E. Coomassie dye-stained SDS–PAGE gel of purified recombinant Crn1 protein variants.
F. High-speed actin cosedimentation assays of Crn1-FL and truncated Crn1 variants in F-buffer. F-actin-bound coronin (µM/µM) was plotted against free coronin (µM) (data from three technical replicates for each protein variant) and fitted using the Hill equation.
G. Representative negatively stained transmission electron microscopy (TEM) images of preassembled F-actin (2 µM) incubated with Crn1 variants at 1 µM in F-buffer. Scale bar, 50 nm.

To understand the mechanisms underlying IDR function, we then obtained the recombinant Crn1-FL protein and the same set of truncated variants (Figure 1E). First, their interactions with F-actin were tested in a high-speed actin cosedimentation assay. During the titration of each Crn1 variant to a fixed concentration of actin, Crn1-FL showed a high affinity with a K_d_ = 0.24 µM (Figure 1F; Figure 1-figure supplement 1A), consistent with previous reports ^29^. However, the actin-binding activities of Crn1-ΔN and Crn1-ΔCC were much weaker (Figure 1F), consistent with their *in vivo* delocalization from actin patches (Figure 1C). Both Crn1-N and Crn1-CC were found to bind F-actin but their affinity for F-actin was much lower than that of the full-length protein (Figure 1F) ^31,39^. Interestingly, Crn1-IDR did not show apparent affinity toward F-actin (Figure 1F), which suggests that the absence of Crn1-ΔIDR colocalization with actin patches was not due to loss of the interaction between the IDR and F-actin. By analyzing the Crn1-ΔIDR protein via size-exclusion chromatography (SEC) and transmission electron microscopy (TEM), we found that Crn1-ΔIDR displayed a more heterogeneous profile than Crn1-FL in the form of high-order soluble assemblies (Figure 1-figure supplement 1B,C). In high-speed ultracentrifugation, while Crn1-FL and other truncated Crn1 variants remained in the supernatant, approximately half of Crn1-ΔIDR was found in the pellet fraction (Figure 1-figure supplement 1D,E). Such aberrant protein behavior hampered the use of a conventional high-speed actin cosedimentation approach to characterize the binding of Crn1-ΔIDR to F-actin. Hence, we directly compared the abilities of different Crn1 proteins to crosslink with F-actin.

Neither the single domains alone (Crn1-N, Crn1-IDR or Crn1-CC) nor the domain-truncated variants (Crn1-ΔN, Crn1-ΔIDR or Crn1-ΔCC) were able to bundle F-actin like Crn1-FL (Figure 1G; Figure 1-figure supplement 1F). In the presence of F-actin, Crn1-ΔIDR showed heterogeneous ring-like assemblies without a clear association with actin filaments (Figure 1G; Figure 1-figure supplement 1F). The above results suggest the indispensable role of IDR in maintaining the appropriate Crn1 conformation and folding to prevent the formation of dysfunctional self-assemblies.

### IDR regulates Crn1-CC oligomerization

Besides the IDR roles in facilitating folding properties shown above, we also hypothesized that the IDR indirectly regulates actin assembly by modulating the neighboring functional domains of coronin, such as the CC domain, which is known to control coronin oligomerization and function ^26,27,29,31,50^. We first characterized the physico-chemical properties of recombinant Crn1-CC and Crn1-IDR using circular dichroism spectroscopy ^51–53^. The far-UV spectrum of Crn1-IDR showed a large negative ellipticity at 198 nm, indicating disorder, whereas Crn1-CC generated positive ellipticity below 200 nm and two minima at 208 and 222 nm, indicating a helical conformation (Figure 2-figure supplement 1A,B). We then analyzed Crn1-CC using the Deepcoil2 engine ^54^, which assigned a heptad pattern (abcdefg) to the α-helical Crn1-CC ^55–57^. The helical coiled coil property of the C-terminus is a generally conserved feature among different coronin family members, e.g., the CC domain of murine Coronin 1A (Figure 2-figure supplement 1C), which forms a well-defined trimeric CC, as shown by X-ray crystallography and *in vivo* characterizations ^36,43^. We found that the critical ‘a’ and ‘d’ positions of the Crn1-CC ‘heptad’ are mainly occupied by the hydrophobic residues Leu and Val (Figure 2-figure supplement 1D). However, the ‘a’ and ‘d’ positions within one of the heptads are not filled by hydrophobic residues (Figure 2-figure supplement 1D, circled residues). Such heterogeneity prevents the precise prediction of Crn1-CC oligomeric state, either empirically or by relying on CC prediction algorithms ^58^.

To quantitatively determine the oligomerization state of Crn1-FL and its truncated variants, we next performed analytical ultracentrifugation sedimentation velocity (AUC-SV) experiments under physiological ionic strength with 150 mM NaCl. Crn1-FL (Figure 2A, red curve) sedimented as two species with sedimentation coefficients of 3.6 S (MW_app_ = 83 kDa) and 6.5 S (MW_app_ = 286 kDa), consistent with monomers and tetramers, respectively. Deletion of the C-terminal CC domain (Crn1-ΔCC) produced a single band at 3.4 S (MW_app_ = 69.5 kDa) corresponding to a monomeric form (Figure 2A, blue curve). Thus, the tetrameric form of Crn1 requires the CC domain. Next, we asked if the CC domain is sufficient for Crn1 tetramerization. To our surprise, the *c(s)* profile of Crn1-CC showed a single peak at 5.3 S (MW_app_ = ~120 kDa) close to the peak predicted for a pentadecamer (Figure 2B, red curve). In contrast, Crn1-ΔN, which contains both the CC and IDR domains, showed a single species at 3.4 S with an MW_app_ of ~116 kDa, consistent with a tetramer (Figure 2B, blue curve). Such results suggest that although the IDR domain alone is a monomer (Figure 2-figure supplement 1E), it is critical in maintaining the tetrameric state of the adjacent CC domain, which otherwise forms higher-order oligomers.

**Figure 2.**
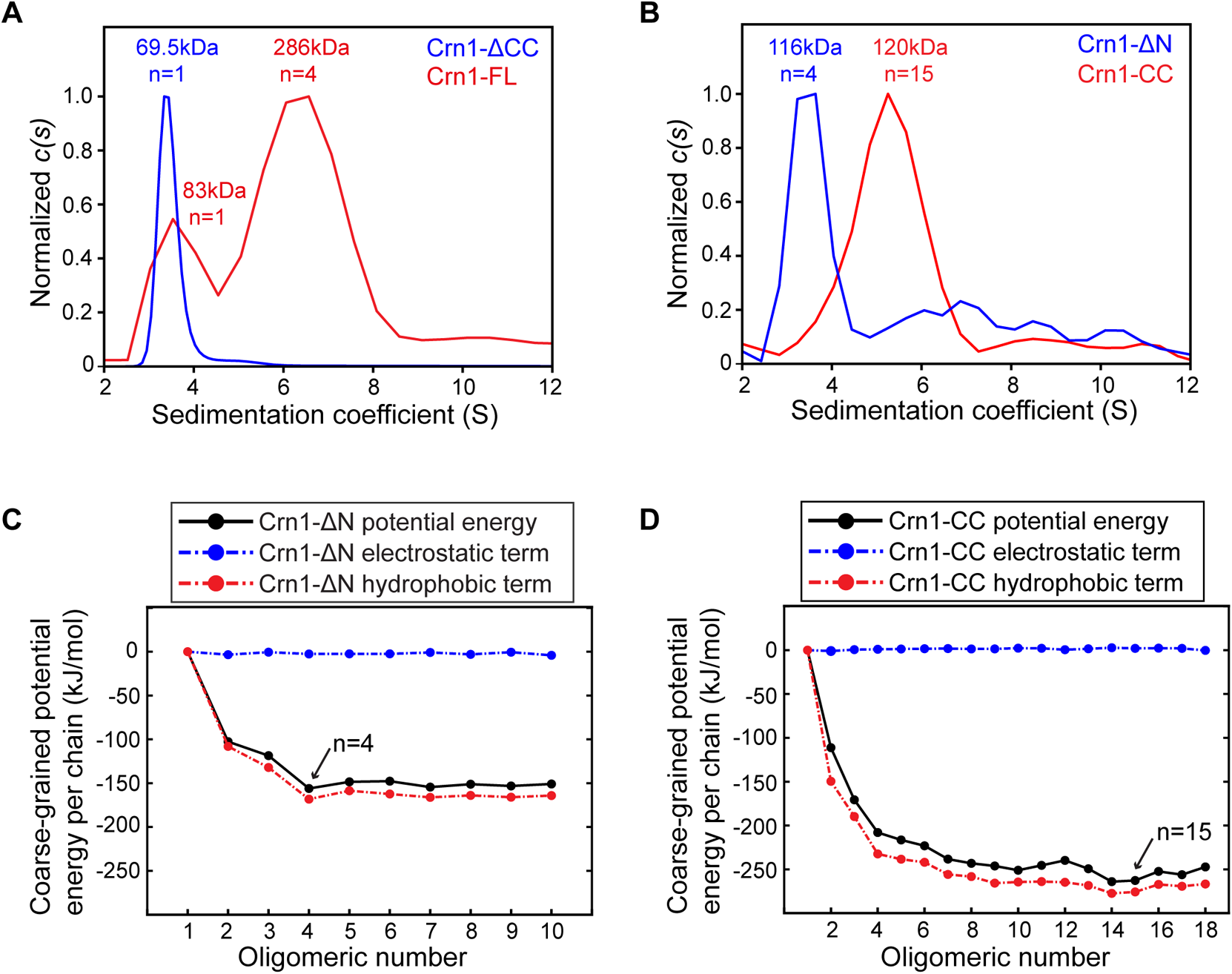
IDR regulates Crn1-CC oligomerization. (A-B) Overlaid AUC-SV profiles of Crn1 protein variants. Sedimentation coefficient distribution *c(s)* profiles of Crn1-FL (8.5 µM) and Crn1-ΔCC (9.2 µM) (A) and Crn1-ΔN (4.25 µM) and Crn1-CC (3 µM) (B). The *c(s)* distribution was normalized to max *c(s)* in GUSSI. Estimated molecular weights and oligomeric states are indicated.
(C-D) Potential energy per chain of Crn1-ΔN (C) and Crn1-CC (D) varies with the number of helices in coarse-grained simulations. Potential energy was decomposed into a hydrophobic term and an electrostatic term for analysis of their contributions.

The IDR has been reported to modulate its own hydration-free energy and that of the connected folded domain, which might tune the hydrophilicity of the protein as well as macromolecular activity ^59^. We were then motivated to ask if the IDR can tune the energy landscape of the neighboring CC domain. We used coarse-grained molecular simulations to investigate the relationship between potential energy per chain and the degree of oligomerization. We simulated Crn1-ΔN and Crn1-CC, the CC domain with and without the IDR, respectively, at the same time. Since knowledge about the arrangement of chains in Crn1-CC oligomers was limited, starting configurations arranged in both parallel and antiparallel alignments were tested, and coarse-grained simulations actually showed indistinguishable results for the two kinds of alignments (Figure 2-figure supplement 1F). Thereafter, we used only the parallel orientation in subsequent simulations. Consistent with the AUC-SV experiments (Figure 2B), the coarse-grained potential energy per chain did not drop further after the tetrameric state was reached for Crn1-ΔN, showing a maximum stabilization effect at the tetrameric state (Figure 2C, black curve). To depict the energy contributions, we carried out energy decomposition and found that hydrophobic interactions showed a dominant effect in guiding the overall potential energy per chain along the examined oligomerization states of Crn1-ΔN (Figure 2C, red curve). This suggests that the hydrophobic residues within the CC domain are important in determining the oligomerization state of Crn1-ΔN. In contrast, electrostatic interactions had a negligible effect on the optimization of Crn1-ΔN oligomerization, as the electrostatic part of the potential energy per chain changed unnoticeably among Crn1-ΔN complexes of different numbers (Figure 2C, blue curve). Surprisingly, without the IDR, the potential energy per chain of Crn1-CC showed a further decrease after the tetrameric state was reached until the tetradecameric-to-pentadecameric state transition, suggesting that Crn1-CC alone tends to form high-order oligomers in the absence of the IDR (Figure 2D), again consistent with the AUC-SV results (Figure 2B). Similar to Crn1-ΔN, hydrophobic interactions, rather than electrostatic interactions, within Crn1-CC were the underlying regulatory factors that determined the optimal oligomeric number, additionally supporting the critical roles of hydrophobic interactions between heptad repeats of Crn1-CC.

### Optimized oligomerization of Crn1 regulates *in vivo* Crn1 functionality

We next asked whether the fine-tuning of oligomerization influences Crn1 functionality in actin assembly, both *in vivo* and *in vitro*. *In vivo* homo-oligomers have been investigated by density gradient centrifugation and gel filtration chromatography ^36,60^. However, it is challenging to quantify the *in vivo* oligomer states for the proteins that are able to undergo tunable heterogenous self-assembly oscillating between different oligomeric orders and often co-sedimented with associated binding partners *in vivo*. In addition, the oligomeric status for tunable oligomers is sensitive to the conditions used for cell lysate preparation. To tackle such a challenge to study in vivo hetero-oligomers quantitatively, we employed a protein engineering approach to determine the oligomerization-status-dependent functions of yeast Crn1, both *in vivo* and *in vitro*. Here, we designed and engineered Crn1 variants at different homo-oligomeric states, ranging from dimer to pentamer, by replacing the Crn1 CC domain with several well-characterized homo-oligomeric CC motifs ^61^ (Figure 3A). *De novo*-designed *CRN1* variants underwent genomic integration into the background of *crn1Δ* yeast under the endogenous promoter and tagged by C-terminal mRuby2, allowing *in vivo* examination of their localization at actin patches. All the engineered homo-oligomeric Crn1 variants were expressed at similar total protein levels *in vivo*, as shown by western blot assay (Figure 3B). Through mRuby2-based fluorescent imaging and image analysis, we found that homo-oligomeric Crn1 variants generally maintained actin patch localization (Figure 3C), indicating an association with F-actin even with different oligomerization motifs ^44^. In addition, by quantifying the signal ratios of Crn1 variants on the actin patch over the total cellular signals (Figure 3-figure supplement 1A), we found that Crn1, Crn1ΔCC-Di and Crn1ΔCC-Tri at lower oligomeric states exhibited lower signal enrichment on actin patches than higher-level oligomeric Crn1 variants, Crn1ΔCC-Tet, and Crn1ΔCC-Pent as well as Crn1 WT (Figure 3D). The latter three variants showed similar actin patch signal ratios, suggesting a similar functional association to F-actin in the patch.

**Figure 3.**
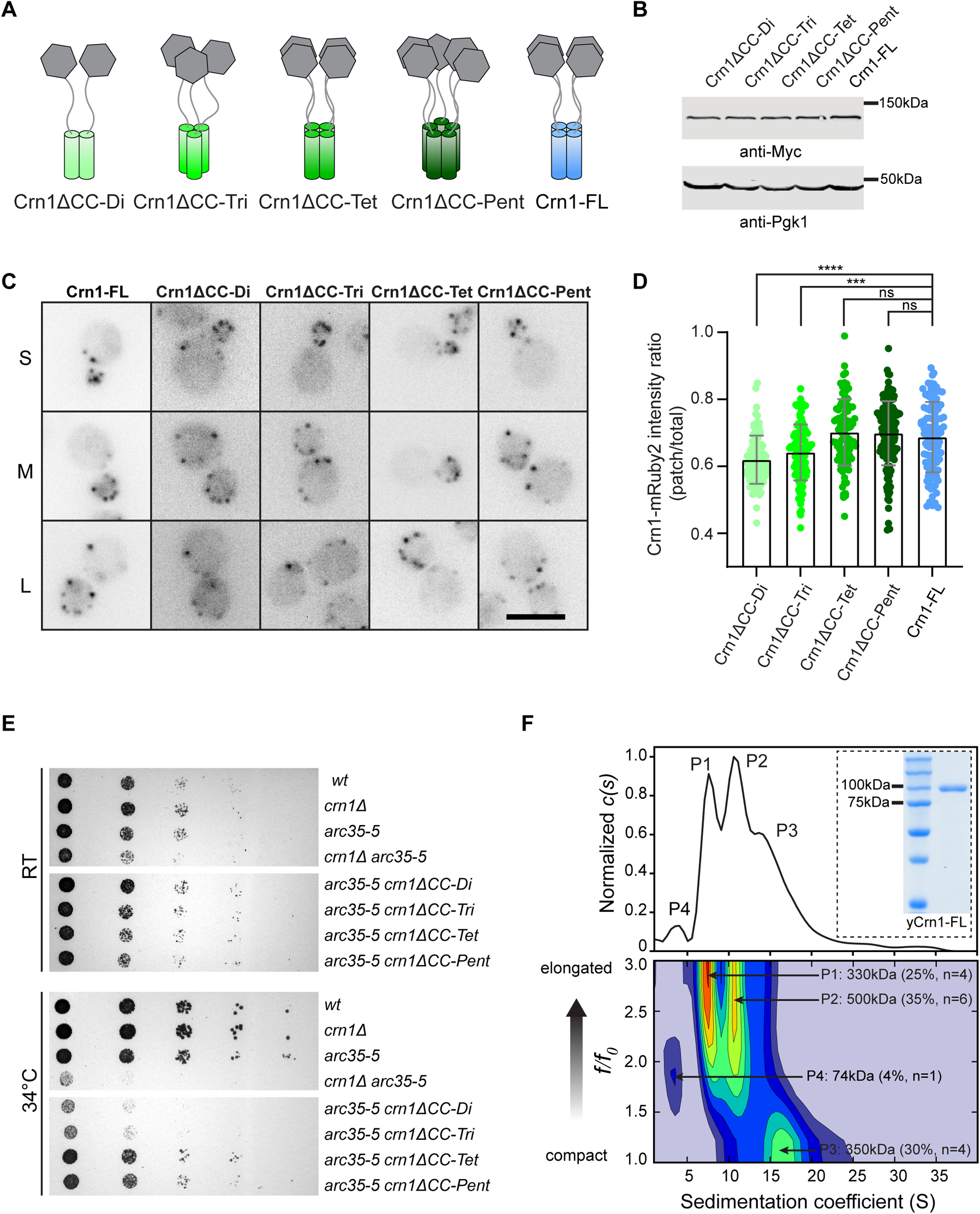
Optimized oligomerization of Crn1 regulates *in vivo* Crn1 functionality. A. Cartoon diagram of *de novo*-designed homo-oligomeric Crn1 variants. *In vivo* yeast expression was through genomic integration with mRuby2 and 4xMyc tag at the C terminus under the native promoter.
B. *In vivo* homo-oligomeric Crn1 variants in (A) examined by western blot using total cell lysate and antibodies against Myc-tag. The anti-Pgk1 antibody was used as a loading control.
C. Representative fluorescent images of mRuby2-tagged homo-oligomeric Crn1 variants in yeasts. S, small-budded cell; M, middle-budded cell; L, large-budded cell. Scale bar, 5µm.
D. Crn1-mRuby2 signal intensity ratio on actin patches (patch /total, see Methods) in (C). From left to right n=98, n=108, n=95, n=149, n=102 patches. Bar graphs indicate mean values. Error bars: SD.
E. Yeast spotting assay of homo-oligomeric Crn1 variants in the genetic background of *crn1Δ arc35-5*. Cells were grown on YPD at the indicated temperature for 42 hours before imaging.
F. Size and shape distribution profile of yeast purified full-length Crn1 (namely: yCrn1-FL) (5.8 µM) as shown in dashed box from AUC-SV experiments performed in 50 mM Tris pH 8.0, 150 mM NaCl buffer and analyzed by *c(s, ff0)* model. Different populations (P1 to P4) shown in *c(s)* profile were analyzed by sedimentation coefficient (S) and frictional ratio *(f/f0),* and generated the heat map in the lower panel. Color in heat map indicates concentration of different populations, from lowest (blue) to highest (red). Increasing value of *f/f0* indicates the elongated or nonglobular species. Unpaired two-tailed Student’s t test assuming equal variance was used to determine difference between 2 groups in (D), *p < 0.05; **p < 0.01; ***p < 0.001; ****p < 0.0001, ns = not significant.

To provide a sensitive system to evaluate the fine-tuning of Crn1 functionality *in vivo,* we compared growth defects due to homo-oligomeric Crn1 variants in the background of *arc35-5,* a temperature-sensitive mutant of the Arp2/3 complex ^26,27^. *arc35-5* exhibits a negative genetic interaction with *crn1Δ*, resulting in lethality at high temperature ^26,27^. At an intermediate temperature of 34°C (semi-permissive for *arc35-5*), the *crn1Δ arc35-5* double mutant showed severe synthetic sickness (Figure 3E). Lack of any individual domain among the IDR, the CC domain, or the N-terminal β-propeller in the background *arc35-5* led to a level of lethality similar to that of *crn1Δ arc35-5* at 34°C (Figure 3-figure supplement 1B). The homo-oligomeric Crn1 variants exhibited different sensitivities in the *arc35-5* background at 34°C. *arc35-5 crn1ΔCC-Di* and *arc35-5 crn1ΔCC-Tri* showed similar severe growth defects comparable to those of *crn1Δ arc35-5*, whereas the growth defect of the *crn1Δ arc35-5* double mutant at 34°C was restored in *arc35-5 crn1ΔCC-Tet* and *arc35-5 crn1ΔCC-Pent* (Figure 3E). Assessments of actin patch localization and genetic interactions indicate that the oligomerization state of Crn1 fine-tunes its functionality *in vivo*. Tetrameric and pentameric Crn1 exhibited better functionality than the dimeric or trimeric Crn1 variants, suggesting the necessity to maintain Crn1 oligomerization for optimal functionality (Figure 2A).

In addition, we sought to provide a biochemical characterization of Crn1 oligomerization status in yeast. We have expressed and purified Crn1 from yeast cells (namely yCrn1-FL) using a previously reported high-yield overexpression system ^62^ (Figure 3F, dashed box) that generated sufficient materials for AUC analysis in a solution of physiological ionic strength of 150 mM NaCL (Figure 3F). As a result, yCrn1-FL mainly displayed two populations of oligomers. 2D size-and-shape analysis characterized that around 55% of yCrn1-FL is present in tetrameric states, whereas an additional >35% of yCrn1-FL is in a hexamer form. Based on the fictional ratio *f/f0* that is related to the particle shape, the tetrameric yCrn1-FL proteins are mixed with elongated (*f/f0* >2) and globular species (*f/f0*~ 1.2) (Figure 3F), whereas bacteria purified Crn1-FL were mainly in elongated (*f/f0* >2) shape (Figure 3-figure supplement 1C). Together, yeast expressed Crn1 also supports a tetrameric state as a dominant population *in vivo*, although the potential functional difference between elongated and globular-shaped tetramers is unclear and worth future investigations.

### Crn1 oligomeric state regulates F-actin crosslinking and inhibition in Arp2/3-mediated actin nucleation *in vitro*

To further quantitatively characterize the biochemical functions of engineered homo-oligomeric Crn1 variants, we first purified the Crn1 homo-oligomeric variants using an *E. coli* expression system (Figure 4-figure supplement 1A) and examined their oligomerization state by AUC-SV. All Crn1 variants displayed the expected oligomeric form, with one band in the *c(s)* plots (Figure 4-figure supplement 1B). Next, we compared their actin-bundling activities via a low-speed actin cosedimentation assay under physiological ionic strength with 150 mM KCl. Interestingly, we found that the Crn1 variants crosslinked F-actin in an oligomerization state-dependent manner. Crn1ΔCC-Tet and Crn1ΔCC-Pent, which are high-level oligomers, demonstrated higher bundling efficacy than Crn1ΔCC-Di and Crn1ΔCC-Tri (Figure 4A; Figure 4-figure supplement 1C). We also directly visualized the bundling of F-actin using total internal reflection microscopy (TIRF) in the presence of different homo-oligomeric Crn1 variants and quantified bundling efficacy by measuring ‘skewness’ ^63,64^. Similar oligomerization-dependent actin crosslinking was observed (Figure 4B and C).

**Figure 4.**
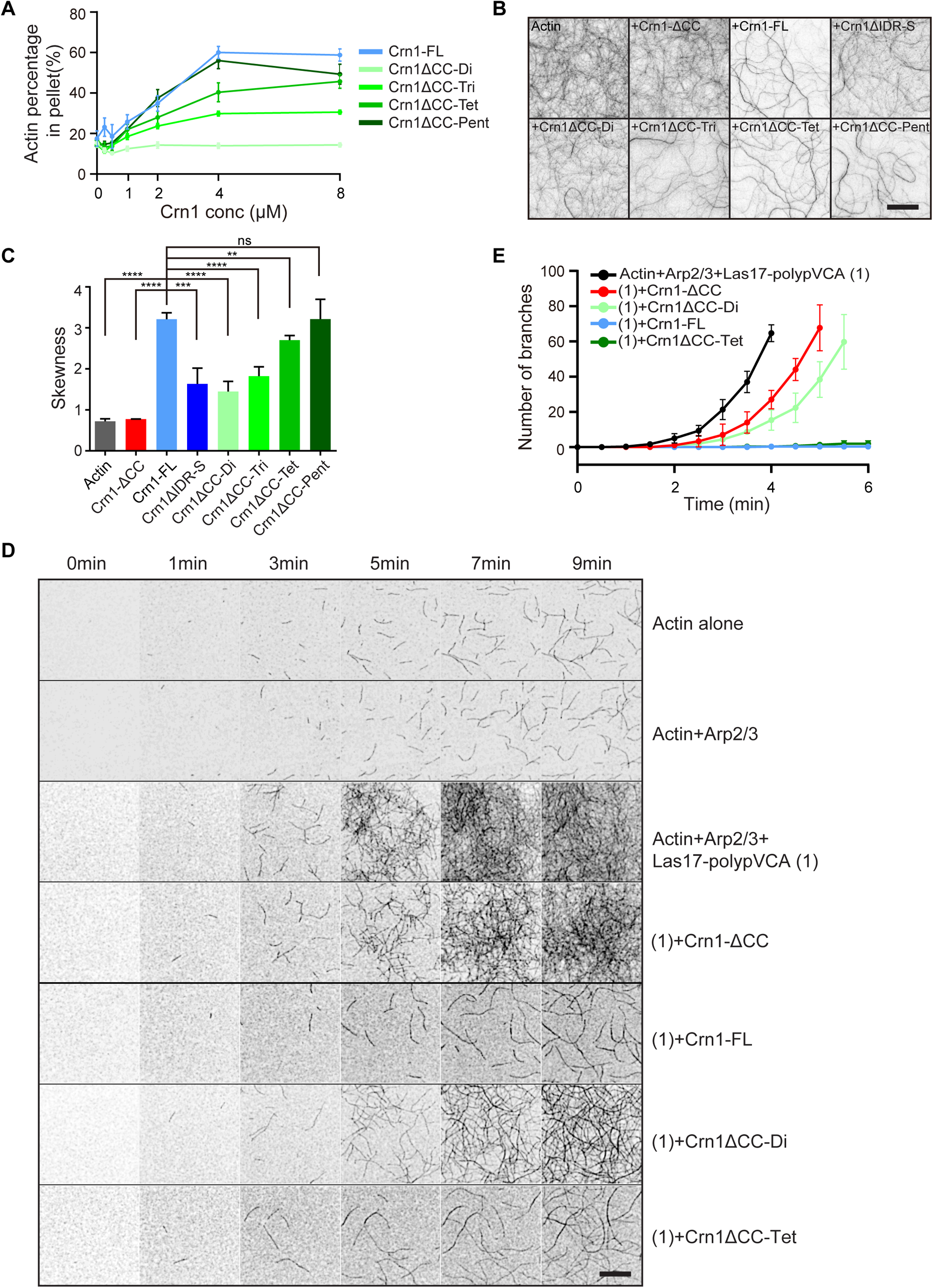
Crn1 oligomeric state regulates F-actin crosslinking and inhibition in Arp2/3-mediated actin nucleation *in vitro*. A. F-actin bundling profile from low-speed actin cosedimentation assay using homo-oligomeric Crn1 variants in buffer with physiological ionic strength of 150 mM KCl. Actin bundling was measured by quantifying actin amount in pellet fraction through densitometry from SDS-PAGE gel using ImageJ, from three technical replicates. Error bars, SD.
B. TIRF images showing Crn1-mediated actin bundling by different Crn1 variants after 90 min. Actin (1 µM) was co-polymerized with Crn1 variants (2 µM) in buffer with physiological ionic strength of 150 mM KCl. Scale Bar: 10µm.
C. Quantification of bundling level in ROI (32×32 µm^2^) from (B) by skewness of pixel intensity. Bar graphs indicate mean values. (n=4 ROIs for each sample, error bars: SD).
D. Representative time-lapse TIRF images of Arp2/3-mediated actin polymerization in the presence of indicated Crn1 variants, including 1 µM actin, 5 nM Arp2/3 complex, 25 nM Las17-polypVCA, and 100 nM Crn1 variants. Scale bar, 10 µm. Corresponding Figure 4-video 1 is available online.
E. Quantification of actin branches number in ROI (32×32 µm^2^) over the first 6 min of TIRF time-lapse imaging in (D). (n=3 ROIs for each condition, error bars, SD). Unpaired two-tailed Student’s t test assuming equal variances was used to determine difference between 2 groups in (C), *p < 0.05; **p < 0.01; ***p < 0.001; ****p < 0.0001, ns = not significant.

We next compared Crn1ΔCC-Di and Crn1ΔCC-Tet, representing the lower- and higher-level oligomeric states, respectively, based on their inhibition of Arp2/3 complex-mediated actin nucleation ^7,8,26^ using a TIRF-based actin polymerization assay. We quantitatively examined nucleation promoting factor (NPF)-Arp2/3-mediated actin nucleation and branching over time in the presence of different Crn1 variants (Figure 4D and E; Figure 4-video 1). The functional core of Las17/WASp, including the polyproline and VCA (polypVCA) domain ^65^, was expressed and purified from *E. coli* (Figure 4-figure supplement 1D). WT Crn1 (Crn1-FL) showed apparent inhibition of both actin nucleation and branching on the Arp2/3 complex, resulting in decreased overall actin polymerization (Figure 4D and E), consistent with previous work ^7,8,26^. However, deletion of the CC domain (Crn1-ΔCC) did not inhibit the Arp2/3 complex, underscoring the importance of Crn1 oligomerization in inhibiting the NPF-Arp2/3 complex (Figure 4D and E). These results are also consistent with findings from previous pyrene-actin assays ^26^. Under the same experimental conditions, dimeric (Crn1ΔCC-Di) and tetrameric (Crn1ΔCC-Tet) Crn1 variants impaired polypVCA-Arp2/3 complex-mediated actin polymerization to different degrees. Crn1ΔCC-Tet behaved like Crn1-FL, effectively inhibiting Arp2/3-polyp-VCA complex-mediated actin branching (Figure 4D and E). In contrast, Crn1ΔCC-Di exerted a much lower inhibitory effect. Compared with Crn1-ΔCC, however, Crn1ΔCC-Di still had a slight negative impact on Arp2/3-NPF activities (Figure 4D and E). This indicates that Crn1ΔCC-Di was still partly functional, consistent with its partial defect in actin patch localization and partial rescue of the genetic sickness of the *crn1Δ arc35-5* double mutant (Figure 3D and E). Taken together, the aforementioned *in vivo* genetic and cell biology results and *in vitro* actin biochemical experiments suggest that Crn1 function depends on its oligomeric state. Among the range of oligomerization states we examined here, tetrameric Crn1 demonstrated better functionality than lower-level Crn1 oligomers.

### Evolutionarily conservation of IDR in coronin family proteins

The function of the IDR in fine-tuning CC domain oligomerization motivated us to ask whether and how this mechanism is evolutionarily conserved. We analyzed the IDR at the N-terminus of the CC domain across coronin family proteins. We collected 392 coronin homologs from 200 species, including species from all major phyla, based on a previously reported phylogenetic analysis of coronin proteins ^66^, and generated a taxonomic tree from these homologs (Figure 5). We defined their IDR by identifying each unique region between the conserved β-propeller and CC domains. Each IDR was also analyzed by IUPs (website: https://iupred2a.elte.hu/) to validate low-complexity sequences. We found that the presence of the IDR is evolutionarily conserved among coronin family proteins, although the IDR is conserved in neither sequence nor in length, which is a common feature of the IDR ^67,68^. Coronins in only three out of 200 species, *T.vaginalis* (protozoan), *A.anophagefferens* (protozoan), and *B.natans* (unicellular eukaryotic alga), do not contain an IDR. Then, we manually mapped the analyzed IDR into the coronin taxonomic tree (named IDR-MAP hereafter), in which the IDR length is shown in scale (Figure 5). Strikingly, we found that although the IDRs vary in general, they follow a clustered pattern in which the IDR length within each taxonomic clade is similar but clearly different from that of other clades. The coronins of most of the protozoa (including SAR, Excavata and Amoebozoa) contain a short IDR (< 50 aa), and coronins from only Trypanosomatidae of Excavata and Aconoidasida of Alveolata contain a long IDR (between 50-100 aa or >100 aa). However, in the fungal kingdom, all coronins under Ascomycota (branch marked with a black star), including *S. cerevisiae*, carry a long IDR (> 150 aa). In contrast, coronins in most metazoans, especially mammals, have a short IDR (< 90 aa), except for Nematoda, in which the IDR is generally longer (> 150 aa). Our coronin IDR-MAP suggests a potential evolutionary selection mechanism that determined IDR length.

**Figure 5.**
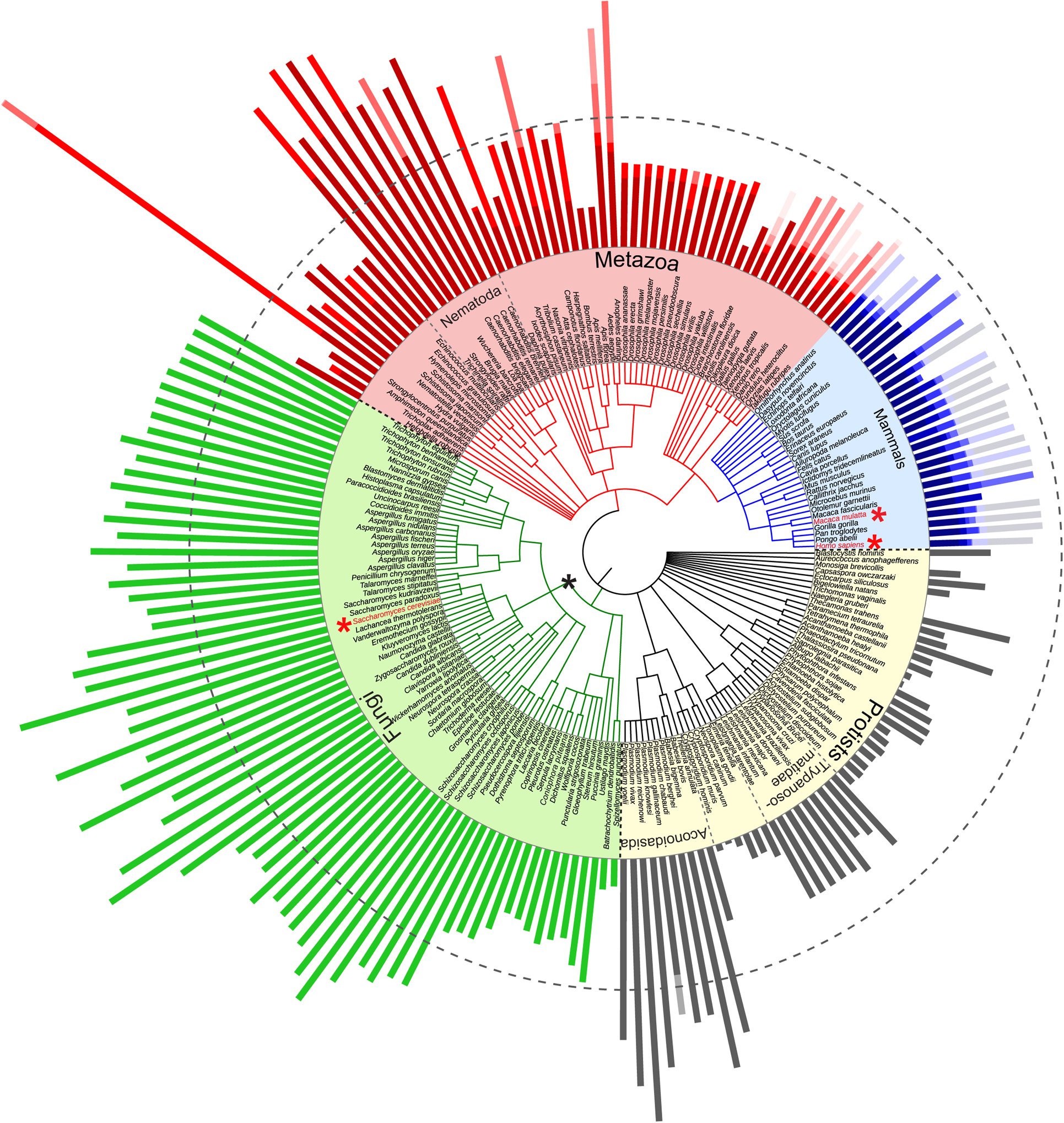
IDR-MAP of coronin family proteins. NCBI Common Taxonomy Tree for 392 coronin homologs from 200 species ^66^. The circular tree was rendered using Interactive Tree Of Life (iTOL) (https://itol.embl.de) ^126^. Three Main parts (protists, fungi and metazoa) were separated by black dashed lines. Bars outside of the tree represent the relative length of coronin IDRs from all analyzed homologs. Additional gradient color bars were overlaid for the species that carry coronin isoforms. The gray dashed circle marked the position of length of 100 aa.

### The evolutionary interplay between IDR length and the stability of CC oligomers

The evolutionarily conserved presence of the IDR in coronin proteins (197 from 200 species) motivated us to ask whether and why the IDR is necessary but varies in length among homologs. We were unable to perform a global investigation of coronin homologs at the same depth. Here, we investigated two well-studied human coronin homologs, Coronin 1A and Coronin 1C ^2,24,36,38^. We compared the *in vivo* localization of full-length Coronin 1A with that of Coronin 1C (1A-FL and 1C-FL) and their respective truncated IDR variants (1A-ΔIDR and 1C-ΔIDR) (Figure 6A) by expressing C-terminal tagged GFP fusion proteins in mammalian cells. Coronin protein patterns and the phalloidin-stained actin cytoskeleton were imaged and analyzed using transiently transfected mouse embryonic fibroblasts (MEFs). The majority of the examined MEFs (> ~60%) showed a high level of F-actin-localized 1A-FL and 1C-FL at cortical filaments or intracellular stress fibers (Figures 6B, D, E; Figure 6-figure supplement 1). However, most (> ~70%) of the 1A-ΔIDR- and 1C-ΔIDR-expressing cells showed a diffuse coronin signal in the cytoplasm without any obvious colocalization with actin filaments (pattern 2) (Figure 6C-E). Such a result suggests the necessity of the coronin IDR in regulating human coronin functions in F-actin association, similar to the IDR-dependent actin association of yeast Crn1.

**Figure 6.**
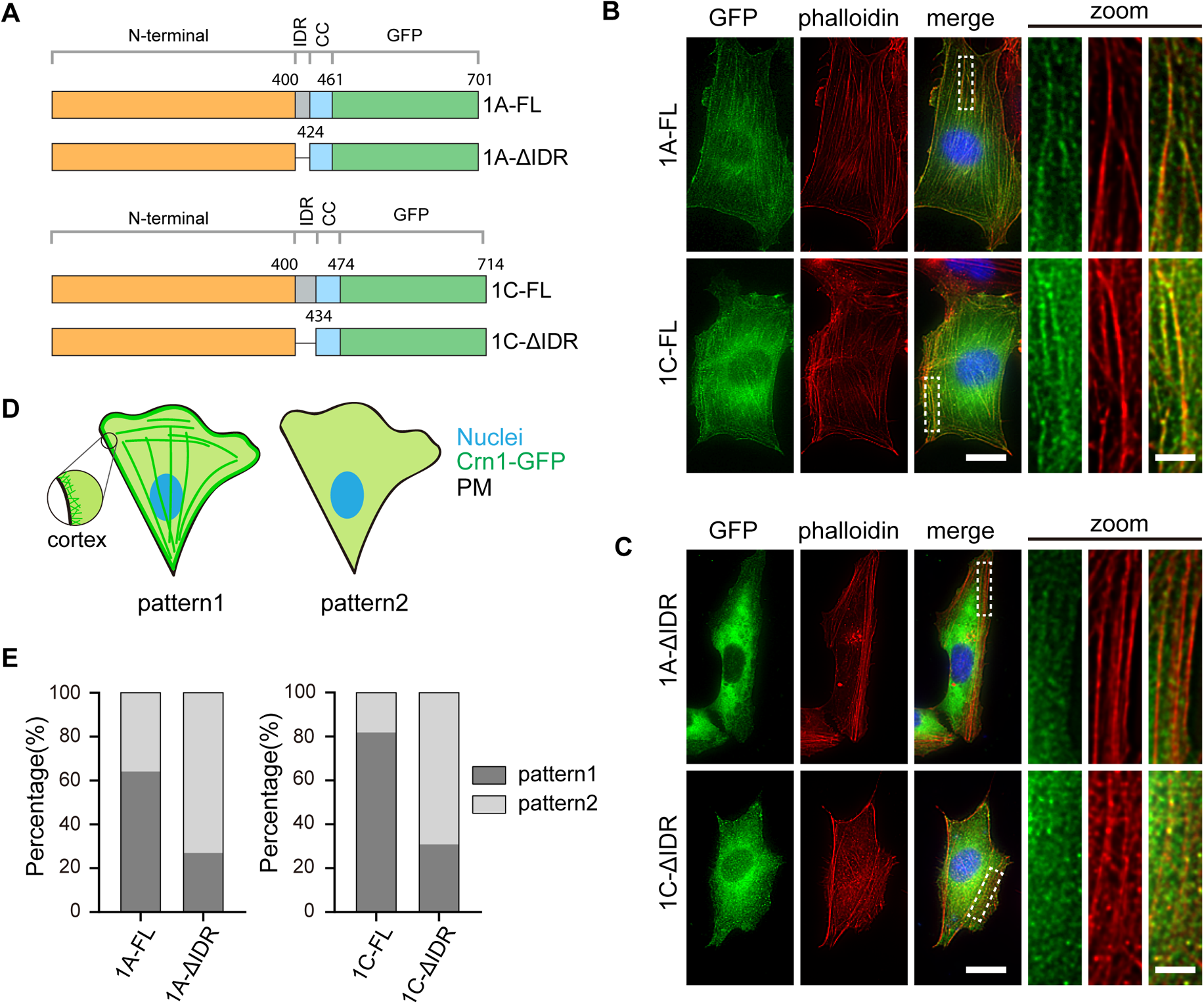
IDR roles in mammalian coronins for F-actin colocalization. (A) Schematic representation of GFP fused Coronin 1A and Coronin 1C constructs for overexpression in mouse embryonic fibroblasts (MEF) cells.
(B-C) Representative images of MEF cell expressing Coronin 1A-GFP and Coronin 1C-GFP with (B) and without IDR (C). Zoomed images were generated from white dashed boxes. Scale bars are 20 µm (left) and 5 µm (zoom).
(D-E) Illustrations of population classification (D) and quantification (E) of coronin-expressing cells based on filamentous and diffused patterns of coronin-GFP observed in (B) and (C). 1A-FL, n=78; 1C-FL, n=33; 1A-ΔIDR, n=52; 1C-ΔIDR, n=55 cells.

The importance of the IDR for human and yeast coronins motivated us to further ask why coronin proteins within distinct evolutionary clades maintain IDRs of different lengths. We first tested whether a shorter IDR would be sufficient for yeast Crn1 functions by maintaining a short, random 40-aa middle fragment of Crn1-IDR (Crn1ΔIDR-S, Figure 7A). Crn1ΔIDR-S was expressed at a protein level similar to that of WT Crn1 (Figure 7-figure supplement 1A, B) *in vivo* but showed decreased localization on actin patches (Figure 7-figure supplement 1C). Crn1ΔIDR-S expression did not demonstrate apparent rescue of the genetic sickness of *crn1Δ arc35-5*, indicating the incompetency of Crn1ΔIDR-S to fully recapitulate Crn1-FL functions (Figure 7B). AUC-SV-based characterization showed that recombinant Crn1ΔIDR-S (Figure 7C) was unable to maintain a tetrameric state. Unlike Crn1-FL, Crn1ΔIDR-S was present as a mixture of trimers and hexamers with sedimentation coefficients of 8.2 S and 13 S, corresponding to sizes of 166 and 322 kDa, respectively (Figure 7D). In addition, Crn1ΔIDR-S exhibited compromised F-actin bundling, as shown by a low-speed actin cosedimentation assay in buffer with a physiological ionic strength of 150 mM KCl (Figure 7E; Figure 7-figure supplement 1D) and TIRF microscopy (Figure 4B and C, Crn1ΔIDR-S). Crn1ΔIDR-S also had a weaker inhibitory effect than Crn1-FL on polypVCA-Arp2/3-mediated actin nucleation and branching (Figures 7F; Figure 7-figure supplement 1E; Figure 7-video 1), consistent with the fact that its oligomerization state is different from that of Crn1-FL (Figures 7D; Figure 2A). We also examined whether the short IDR of Crn1ΔIDR-S would be sufficient to modulate the oligomerization state of Crn1-CC. We applied the same computational approach used for Crn1-ΔN (Figure 2C) and Crn1-CC (Figure 2D) to Crn1-S-IDR-CC. The results showed that the potential energy per chain decreased as the oligomeric number increased in a hydrophobic interaction-dependent manner throughout the scope of investigation (Figure 7G). The fact that the potential energy per chain decreased further after the tetrameric state was reached suggests that a short IDR lacks the capability to stabilize the CC domain in a tetrameric state, as Crn1-FL can (Figure 2C).

**Figure 7.**
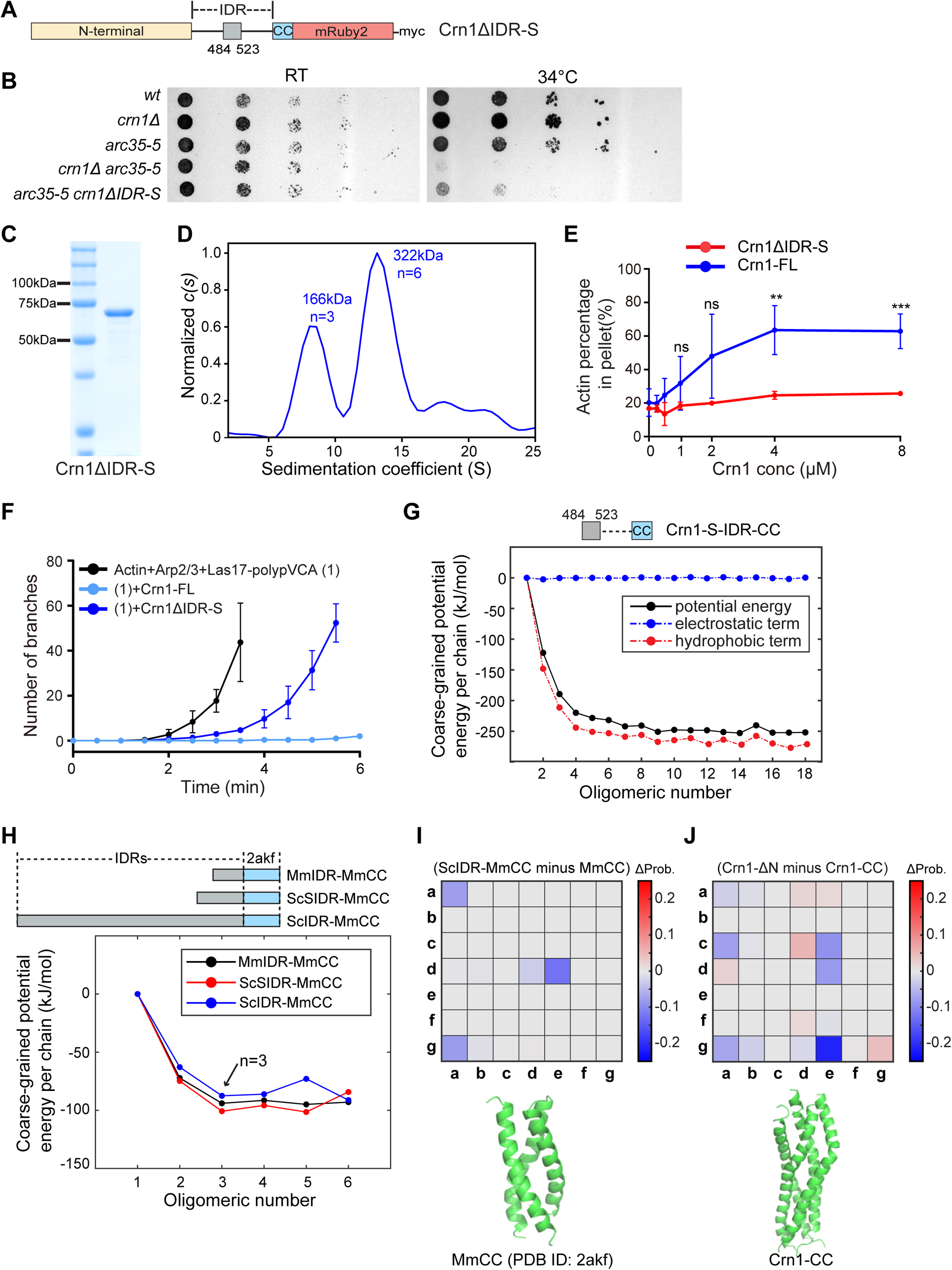
IDR length fine-tunes Crn1 oligomerization state and function. A. Schematic representation of Crn1ΔIDR-S, which was genomically integrated into *crn1Δ* background with C-terminal mRuby2 and 4xMyc tags under the native promoter.
B. Yeast spotting assay of *CRN1ΔIDR-S* in the genetic background of *crn1Δ arc35-5*. Cells were grown on YPD at the indicated temperature for 42 hours before imaging
C. Coomassie dye-stained SDS-PAGE gel of purified recombinant Crn1ΔIDR-S.
D. Sedimentation coefficient distribution profile of Crn1ΔIDR-S (8.1 µM) in AUC-SV. The *c(s)* distribution was normalized to max *c(s)* in GUSSI, estimated molecular weights and oligomeric states are indicated.
E. Bundling profiles of Crn1ΔIDR-S (n=4 from two biological replicates with two technical replicates) and compared to Crn1-FL (n=3 biological replicates) in the buffer with physiological ionic strength of 150 mM KCl from low-speed actin cosedimentation assay. Coronin-induced actin-bundling was measured by quantifying actin amount in pellet fraction through densitometry from SDS-PAGE gel. Error bars, SD.
F. Quantification of F-actin branching events in ROI (32×32 µm^2^) over the first 6 min from TIRF movie. Arp2/3-mediated actin polymerization reaction contained 200 nM Crn1-FL or Crn1ΔIDR-S. (n=3 ROIs for each condition, error bars, SD). See also Figure 7-video 1.
G. Potential energy per chain of Crn1-S-IDR-CC varies with number of helices in coarse-grained simulations. Potential energy was decomposed into hydrophobic term and electrostatic term for analysis of contribution.
H. Potential energy per chain of three recombinants of MmCC with the IDRs, denoted as MmIDR-MmCC (black), ScSIDR-MmCC (red), and ScIDR-MmCC (blue), in coarse-grained simulations. MmCC reprsents murine Coronin1A CC (PDB ID: 2akf). MmIDR-MmCC represents MmCC linking to its native IDR denoted as MmIDR (403-429 aa). ScSIDR-MmCC represents MmCC linking to yeast Crn1 short IDR, denoted as ScSIDR (484-523 aa). ScIDR-MmCC represents MmCC linking to yeast Crn1 full-length IDR denoted as ScIDR (401-604 aa).
I. Motif contact difference map between MmCC and ScIDR-MmCC (top) and a representative side view of MmCC trimer (bottom). Red and blue in motif contact difference map indicate increase and decrease in contact probability, respectively. The representative side view was adapted from its crystal structure (PDB ID: 2akf).
J. Motif contact difference map between Crn1-CC and Crn1-ΔN (top) and a representative side view of Crn1-CC tetramer (bottom). Red and blue in motif contact difference map indicate increase and decrease in contact probability, respectively. The representative side view was adapted from the all-atom structure reconstructed from the representative structure in coarse-grained simulations by PULCHRA ^127^ followed by energy minimization. Unpaired two-tailed Student’s t test assuming equal variances was used to determine difference between 2 groups in (E), *p < 0.05; **p < 0.01; ***p < 0.001; ****p < 0.0001, ns = not significant.

We next sought to understand how the IDR tunes the interaction between adjacent helical chains and consequently modulates the CC domain into optimal oligomeric states. First, using coarse-grained simulations, we investigated how different IDRs influence the oligomerization of the mouse Coronin 1A CC domain (named MmCC), which forms a stable parallel homotrimer in crystals and in solution (PDB ID: 2akf) ^43^. We compared the energy landscape of MmCC when fused with its own IDR (aa 403-429, MmIDR-MmCC), short Crn1-S-IDR (aa 484-523, ScSIDR-MmCC), and Crn1-IDR (aa 401-604, ScIDR-MmCC) (Figure 7H). Intriguingly, all three different recombinants showed no further decrease in energy after the trimeric state was reached, indicating that none of the three IDRs altered the oligomeric state of MmCC (Figures 7H; Figure 7-figure supplement 2A-C), consistent with an X-ray crystallographic study ^43^. Such results suggest that the length of the IDR does not influence the oligomerization of well-packed CC domains, such as the mouse Coronin 1A CC domain (MmCC).

Next, the different roles of the IDR in regulating Crn1-CC and MmCC motivated us to evaluate how different interactions between the helical chains of MmCC are from those of Crn1-CC using the protein structures from simulated tetrameric Crn1-CC and the crystal structure of MmCC. Previous studies have reported that interactions between heptad position pairs ‘a’-‘a’ and ‘d’-‘d’ constitute a hydrophobic core, while ‘e’-‘g’ forms a salt bridge ^69–71^. We used the motif-averaged contact map to describe the interaction strength between two adjacent helices of tetrameric Crn1-CC and trimeric MmCC (Figure 7-figure supplement 2D,G; Figure 7-figure supplement 3A). Between the MmCC helices, the heptad position pairs in which strong interactions occurred were ‘a’-‘a’, ‘a’-‘d’, ‘d’-‘d’ and ‘e’-‘d’, matching the pattern of the hydrophobic core (Figure 7-figure supplement 2D, MmCC). However, Crn1-CC displayed a varying interaction pattern, including many other heptad-position pairs with low-contact probability, suggesting relatively unstable helical packing compared to that of MmCC (Figure 7-figure supplement 2G, Crn1-CC). Interestingly, when different IDRs were linked to MmCC, the pattern and strength of the interactions between MmCC helices were not much altered for any of the three types of IDRs (Figures 7I; Figure 7-figure supplement 2D-F). For example, the motif-averaged contact map of MmCC with a long Crn1 IDR (ScIDR-MmCC) was slightly changed compared to that of MmCC alone (Figure 7I; Figure 7-figure supplement 2D). This difference was expressed as the Euclidean distance, which was 0.17 between ScIDR-MmCC and MmCC (Figure 7I). In contrast, with Crn1-IDR, Crn1-CC showed a much greater change in pre-existing interactions, resulting in a Euclidean distance of 0.28 without introducing any heptad position pairs with new interactions (Figures 7J; Figure 7-figure supplement 2G). The above analysis and experimental results demonstrated that the IDR modulates the packing of Crn1-CC helices, which are unstable due to nonideal interacting residues, but has a less pronounced influence on the well-packed MmCC. Such evolved CC domain sequences and packing patterns of coronin family proteins may thereby contribute to evolutionary selection of IDR length.

## Discussion

### Oligomerization regulates the biochemical activities of ABPs in actin assembly

The initiation, reorganization, and depolymerization of the actin cytoskeleton are biochemically orchestrated by a diverse array of ABPs, including actin nucleation proteins (e.g., Arp2/3 complex, formin), G-ABPs (e.g., profilin), elongation-promoting factor (Ena/VASP), crossing proteins (e.g., fimbrin), and actin depolymerization proteins (e.g., ADF, cofilin) ^72–74^. Emerging evidence shows that inter- or intramolecular interactions of ABPs, including homo- and heterooligomeric interactions and lower-order to higher-order molecular assembly, are fundamental mechanisms in dynamic regulation of the biochemical activities of ABPs, such as during signal transduction, and thereby actin cytoskeleton polymerization and organization ^63,75–97^. For example, during plant immune activation, intermolecular interactions between plant formin dimers on the plasma membrane enhance formin activities in actin nucleation, which plays a critical role in remodeling the actin cytoskeleton of plant cells upon bacterial infection ^63,87,93^. In mammals, multivalent interactions between WASP and the Arp2/3 complex also activate actin nucleation during T-cell signal transduction ^94–96^. Mammalian Ena and VASP are also known to cluster as tetrameters to enhance actin elongation activity by creating a tetravalent G-actin-binding (GAB) domain through a CC domain ^78–80^. After nucleation, F-actin stabilization also requires lower-order protein oligomerization, such as CC-mediated dimeric interactions between tropomyosins, which stabilize F-actin and exclude the binding of other ABPs to F-actin ^81–83^. In addition, dimerization of vinculin through its C-terminal tail is critical for F-actin filament bundling and engages multiple binding partners at focal adhesion sites ^84–86,97^. This oligomerization state-dependent regulation of ABPs through interactions with different binding partners in an oligomerization state-dependent manner suggests sophisticated regulatory mechanisms underlying actin remodeling. The coronin C-terminal domain is predicted to form an oligomeric α-helical CC ^10,36,38, 40–42,60^. Here, we reported the importance of coronin protein oligomerization for fine-tuning its inhibition in Arp2/3 complex-mediated actin nucleation and Crn1-mediated actin bundling, in which the optimal tetrameric state of yeast Crn1 was maintained through coupling the CC domain with an IDR. Because it lacks a well-defined hydrophobic residue pairing pattern within the heptad repeats ^69–71,98,99^, Crn1-CC exhibited a heterogenic higher-order oligomeric state (n> 10) on its own, but Crn1-CC could be optimized by the IDR to finetune the oligomerization status towards a better functional tetramer. Precise characterization of protein oligomer status in vivo is technically challenging.

### IDR tunes the higher-order assembly and functionality of coronins with evolutionary selection

Based on the number of N-terminal β-propeller domain, coronins were initially divided into short and long coronins ^5^. Short coronins are subdivided into type I (e.g., Coronin 1A, 1B and 1C) and type II (e.g., Coronin 2A and Coronin 2B) coronins in metazoans and an ‘unclassified’ class in nonmetazoans (e.g., Crn1 of *S. cerevisiae*, coronin of Dictyostelium) ^66^, whereas long coronins are also classified as type III (e.g., POD1) coronins. Short coronins are universal, and coronin homologs contain three domains: the N-terminal structural β-propeller domain, which contains five WD-repeat motifs ^100–102^; the C-terminal CC domain ^10,36,38, 40–42,60^; and a region between the β-propeller domain and CC domain that was named the “unique region (UR)” due to its large variations in length and sequence along evolution, the signatures of the IDR ^103^. While the IDRs of *S. cerevisiae* Crn1 and *D. melanogaster* Dpod1 have shown microtubule-binding abilities ^29,44,104^, other functions of the IDR remain poorly understood. Here, we carried out a focused analysis of 392 short coronins from 200 species. Our results revealed a previously unknown function of the IDR, which maintains the optimal tetrameric state of Crn1 by suppressing higher-order CC domain oligomerization. IDR-containing proteins (IDPs) work differently than fully structured proteins (e.g., molecular recognition and assembly, protein modification, and spacing chains) ^105^. With multivalent interactions, some IDRs can regulate homo- and heterooligomeric interactions, particularly for dynamic ensembles of macromolecular assemblies ^106,107^. We have recently summarized a unique feature of the IDR of endocytosis proteins, in which we observed a striking correlation between IDR length and temporally regulated protein recruitment for endocytosis progression. Endocytic proteins that arrived earlier have a longer IDR than the later actin polymerization- and scission-related proteins ^103^, suggesting potential evolutionary selection of IDR length depending on functional necessity. Recently, mammalian Eps15 and Fcho1/2, early proteins that arrive at the endocytic site, were found to undergo liquid phase separation through weak interactions to optimize endocytosis initiation due to the presence of long IDRs ^108^. However, the functions of IDRs are diverse and do not necessary facilitate higher-order assembly. For example, certain IDRs may function only through conformational switching between static and disorder states ^109–111^. Several ABPs are generally well folded and contain relatively shorter IDRs or loops, such as the capping protein fimbrin, the Arp2/3 complex, and cofilin ^103^. A 27-aa IDR of yeast Sac6 regulates conformational flexibility between the N-terminal EF-hand domain and actin-binding domain 1 (ABD1) in a phosphorylation-dependent manner ^112,113^. Although the IDRs of fimbrin homologs vary in sequence and length, the presence of an IDR and phosphorylation site is evolutionarily conserved among eukaryotic species ^112^. Here, we report another previously unknown mechanism of IDR in yeast Crn1, which optimizes CC domain-mediated oligomerization to maintain appropriate Crn1 function in F-actin crosslinking and inhibition of the Arp2/3 complex. Using *de novo*-engineered Crn1 homo-oligomers spanning from a dimer to a pentamer, both genetic interaction experiments *in vivo* and actin-based biochemical experiments *in vitro*, including F-actin crosslinking and inhibition of Arp2/3 complex nucleation, demonstrated that the functions of Crn1 in the tetrameric state are optimal.

The CC is a common motif in many IDPs, and IDRs are often located next to the protein oligomerization domain, such as CC or prion-prone domains ^114–116^. The CC domain contains multiple seven-residue heptad motifs and packs into parallel or antiparallel helical bundles through hydrophobic interactions ^57,69,70^. The stability of α-helix assembly is the balanced result of interhelical interactions and depends on multiple determinants, including hydrophobic residues at the ‘a’ and ‘d’ positions, which constitute the hydrophobic core. In addition, diverse packing geometries that do not necessarily follow classic heptad patterns exist ^69–71,117–125^. We observed that the hydrophobic interaction between coronin CC helices is the dominant factor that contributes to the interhelical interactions and CC domain oligomerization. Crn1-CC contains two non-hydrophobic residues at the ‘a’ and ‘d’ positions that generate nonideal heptad repeats (Figure 2-figure supplement 1D). This may lead to unstable packing between helices with multiple relatively weak and heterogenic interactions between heptad repeats (Figures 7J; Figure 7-figure supplement 2G). A neighboring long IDR can modulate the energy landscape of helix packing to prevent higher-order assemblies. Most likely, coronins in higher species (e.g., mouse, human), in contrast, have evolved ideal hydrophobic residues within the heptad, which exhibit robust interhelical interactions without needing a long IDR to maintain an optimal energy level for a low-level oligomeric state. The evolutionarily correlated selection of IDR length in the coronin protein family is striking. For an ideal helical assembly such as the CC domain of murine coronin1A, which contains optimal interacting heptad pairs, length is not critical for modulating the energy landscape of the CC domain. In contrast, the situation with nonideal helical assemblies, such as Crn1-CC, is different. The profound evolution of coronin IDRs with varied lengths is partially driven by how well the neighboring helical chains are packed. Of course, exhaustive tests could not be performed. Therefore, we cannot exclude other factors, such as amino acid composition, that may contribute to evolutionary selection of the IDR based on its role in regulating CC domain oligomerization. Nevertheless, the IDRs of yeast Crn1, human Coronin 1A, and Coronin 1C are indispensable for their functional association with and regulation of the actin cytoskeleton, which at least provides conformational flexibility between the N-terminal β-propeller and C-terminal CC domains. Our future endeavors will focus on identifying the diverse regulatory mechanisms of the IDR, which are molecular grammar- and sequence-dependent.

## Acknowledgment

We thank the National Supercomputing Center Singapore (https://www.nscc.sg/) for assisting the computational work. This study was supported by NTU startup grant (M4081533) and Singapore Ministry of Education (MOE) Tier 3 (MOE2019-T3-1-012) to Y.M.; MOE Tier 1 (2018-T1-001-096) to L.L.; MOE Tier 1 (RT13/19) to J. T; MOE Tier 3 (MOE2019-T3-1-012) to L. N.

## Author contributions

X.H. and Y.M. conceived and designed this study. X.H. performed most of the wet lab experiments. Q.M. contributed to yeast genetic and imaging experiments. W.S. and X.H. performed the AUC experiments with guidance from J.T. Z.H. and F.Z. performed simulation work under the guidance of L.L. and L.N. The manuscript was drafted by X.H. and Y.M. and received feedback from all authors. Y.M. supervised these studies.

## Declaration of interests

The authors declare no competing interests.

## Figure titles and legends

**Figure 4-video 1. TIRF movie of Arp2/3-mediated actin polymerization in the presence of Crn1 homo-oligomer variants.**

TIRF movie of Arp2/3-mediated actin polymerization in the presence of indicated Crn1 variants within time range between 0 to 9 min. For each sample, red arrowheads mark branching events within the time period of Figure 4E. Reaction contains 1 µM actin, 5 nM Arp2/3 complex, 25 nM Las17-polypVCA, and 100 nM Crn1 protein variants. Scale bar, 10 µm.

**Figure 7-video 1. TIRF movie of Arp2/3-mediated actin polymerization in the presence of Crn1ΔIDR-S.**

TIRF movie of Arp2/3-mediated actin polymerization in the presence of Crn1ΔIDR-S or Crn1-FL within time range between 0 to 7 min. For each sample, red arrowheads mark branching events within the time period of Figure 7F. Reaction contains 1 µM actin, 5 nM Arp2/3 complex, 25 nM Las17-polypVCA, and 200 nM Crn1 protein variants. Scale bar, 10 µm.

## Figure supplements

Figure 1-figure supplement 1. F-actin binding of recombinant Crn1 variants.

Figure 2–figure supplement 1. Sequence and structural analysis of Crn1-IDR and Crn1-CC.

Figure 3–figure supplement 1. Characterization of Crn1 *in vivo* functionality.

Figure 4–figure supplement 1. AUC verification and in *vitro* functional assays of Crn1 homo-oligomers.

Figure 6–figure supplement 1. IDR roles in mammalian coronins for F-actin colocalization.

Figure 7–figure supplement 1. IDR tunes Crn1 function in a length-dependent manner.

Figure 7–figure supplement 2. Energy landscape of coronin CCs with different IDRs from coarse-rained simulations.

Figure 7–figure supplement 3. Workflow of generating motif-averaged contact map and motif contact difference maps.

## Source data files

**Figure 1-source data 1. IDR is indispensable for Crn1 localization at the actin patch and its interaction with actin filaments.**

Source data folder provides original western blots of Figure 1D, SDS-PAGE gel images of Figure 1E, and original SDS-PAGE gel images for actin-binding affinity analysis.

**Figure 1-figure supplement 1-source data 1. F-actin binding of recombinant Crn1 variants.**

Source data folder provides original SDS-PAGE gel images of Figure 1-figure supplement 1A, TEM images with representative region highlighted of Figure 1-figure supplement 1B, and TEM images with representative region highlighted of Figure 1-figure supplement 1F, which also generated Figure 1G.

**Figure 1-figure supplement 1-source data 2. F-actin binding of recombinant Crn1 variants.**

Source data folder provides original SDS-PAGE gel images of Figure 1-figure supplement 1D, which were also used for the pelleting analysis of Figure 1-figure supplement 1E.

**Figure 3-source data 1. Optimized oligomerization of Crn1 regulates *in vivo* Crn1 functionality.**

Source data folder provides original western blots of Figure 3B, budding yeast image stacks for patch intensity analysis of Figure 3D, original plates scan of Figure 3E, and original SDS-PAGE gel image of purified yCrn1-FL in Figure 3F.

**Figure 3-figure supplement 1-source data 1. Characterization of Crn1 *in vivo* functionality.**

Source data folder provides original plates scan of Figure 3-figure supplement 1B.

**Figure 4-source data 1. Crn1 oligomeric state regulates F-actin crosslinking and inhibition in Arp2/3-mediated actin nucleation *in vitro*.**

Source data folder provides original SDS-PAGE gel images for actin bundling analysis of Figure 4A, stack image of Figure 4B, image stacks of selected ROIs for skewness analysis of Figure 4C, image stacks of Figure 4D, image stacks of selected ROIs for branching analysis of Figure 4E.

**Figure 4-figure supplement 1-source data 1. AUC verification and in vitro functional assays of Crn1 homo-oligomers.**

Source data folder provides original SDS-PAGE gel image of purified Crn1 oligomeric proteins in Figure 4-figure supplement 1A, original SDS-PAGE gel images of Figure 4-figure supplement 1C, and original SDS-PAGE gel image of purified Las17-polypVCA protein of Figure 4-figure supplement 1D.

**Figure 6-source data 1. IDR roles in mammalian coronins for F-actin colocalization.**

Source data folder provides original fluorescent microscope images of Figure 6B and C, and original images for pattern analysis of Figure 6E.

**Figure 6-figure supplement 1-source data 1. IDR roles in mammalian coronins for F-actin colocalization.**

Source data folder provides original fluorescent microscope images of representative pattern 2 population of Coronin 1A-FL and Coronin 1C-FL, as well as original images of vector control representative.

**Figure 7-source data 1. IDR length fine-tunes Crn1 oligomerization state and function.**

Source data folder provides original plates scan of Figure 7B, original SDS-PAGE gel image of purified Crn1ΔIDR-S in Figurae 7C, original SDS-PAGE gel images for actin bundling analysis of Figure 7E, and image stacks of selected ROIs for branching analysis of Figure 7F.

**Figure 7-figure supplement 1-source data 1. IDR tunes Crn1 function in a length-dependent manner.**

Source data folder provides original western blots of Figure 7-figure supplement 1B, budding yeast image stacks for patch intensity analysis of Figure 7-figure supplement 1C, original SDS-PAGE gel images of Figure 7-figure supplement 1D, and image stacks of Figure 7-figure supplement 1E.

## Additional files

Supplementary file 1. Table-S1: List of yeast strains used in this study.

Supplementary file 2. Table-S2: List of plasmids used in this study.

Supplementary file 3. Table-S3: List of primers used in this paper.

Supplementary file 4. Source data tables for generating graph panels and statistical analysis.

## Methods

### KEY RESOURCES TABLE

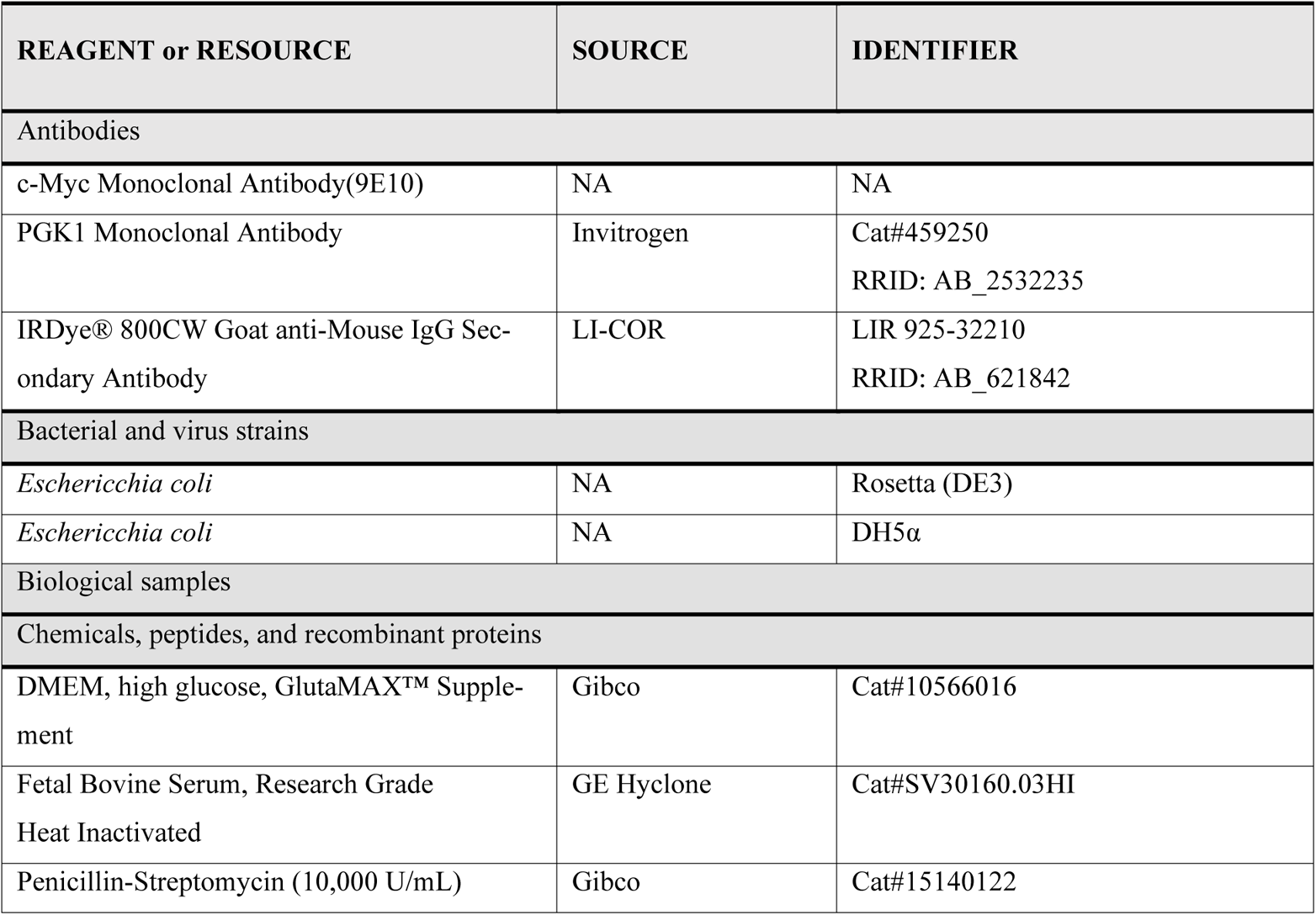
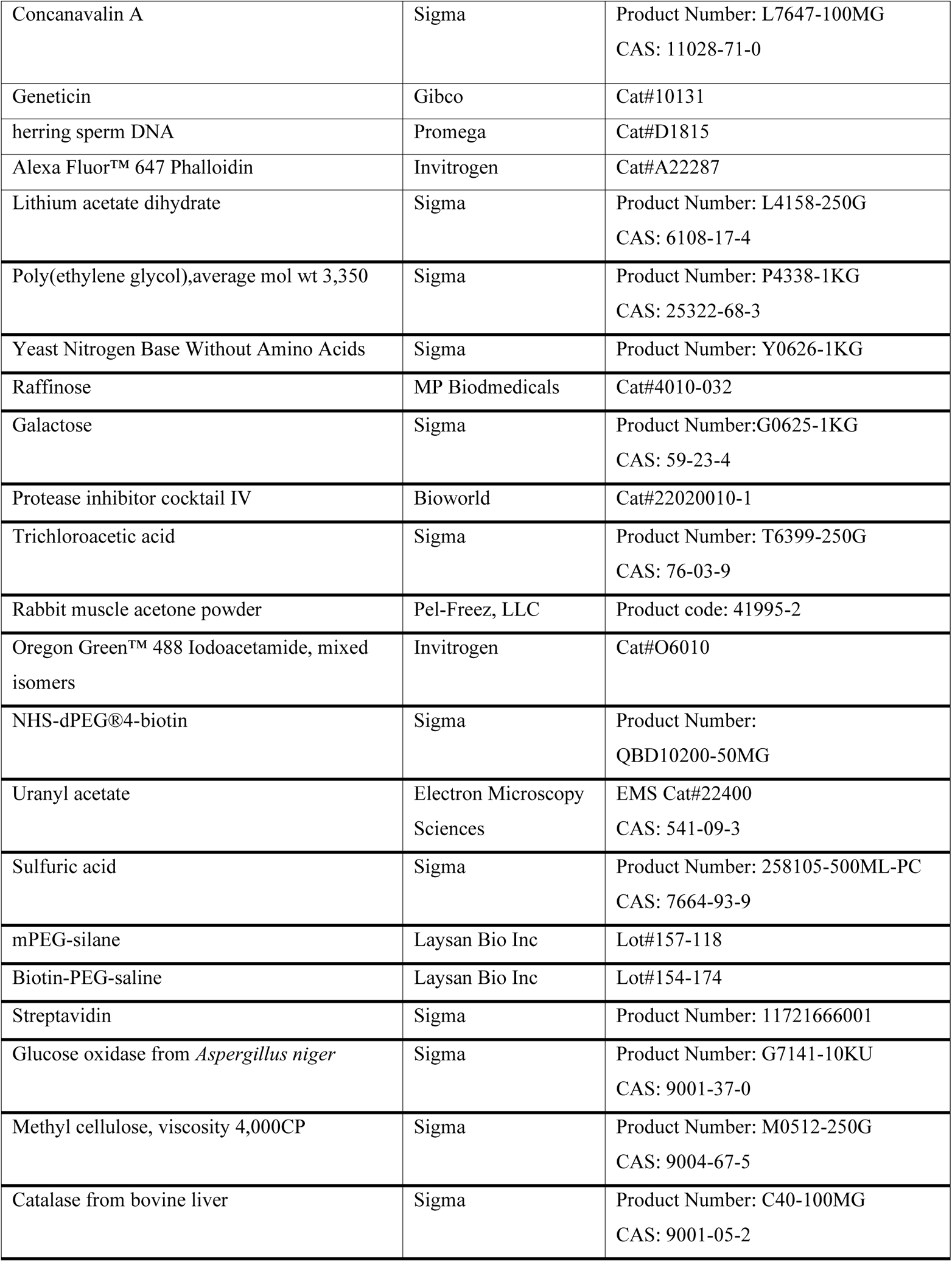
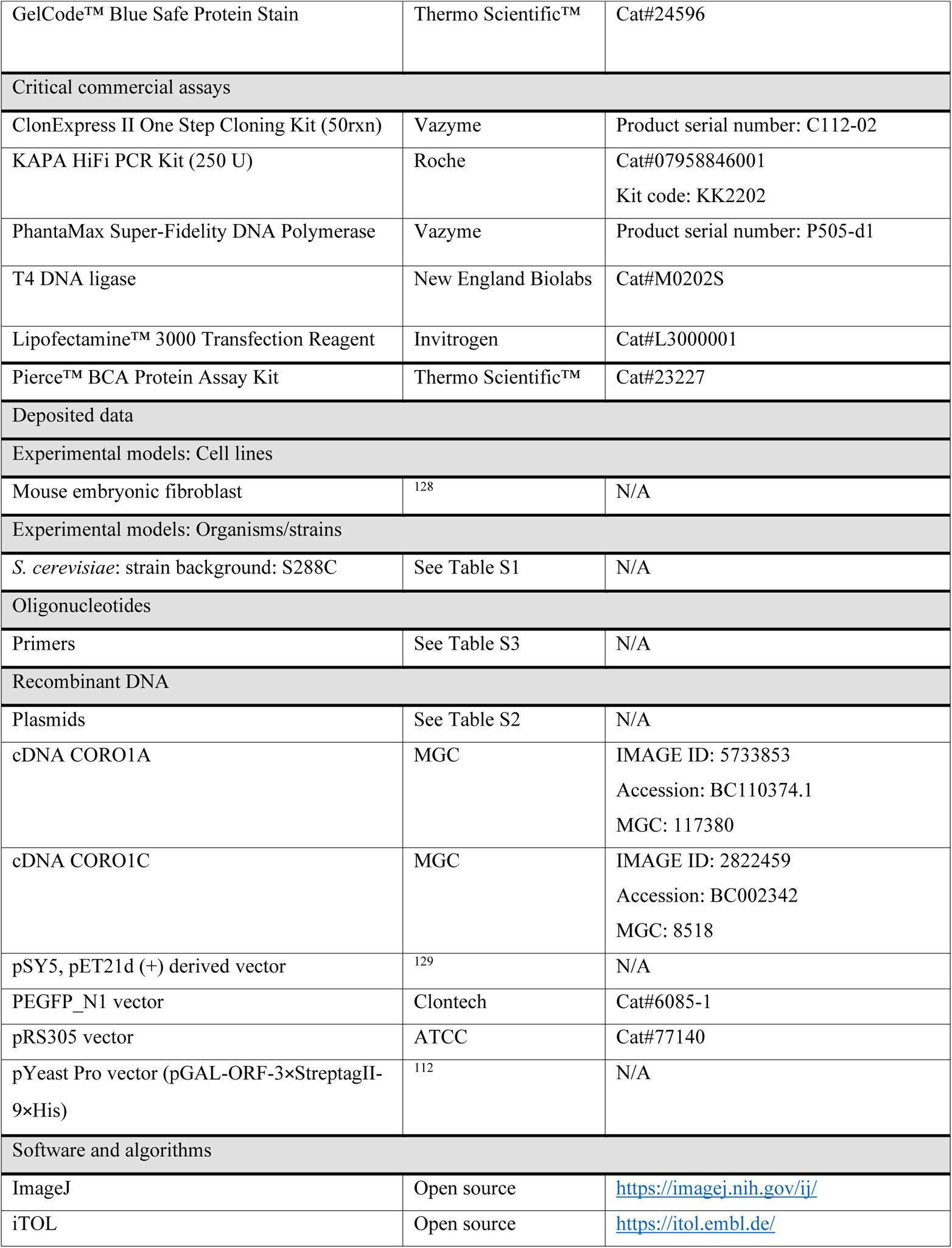
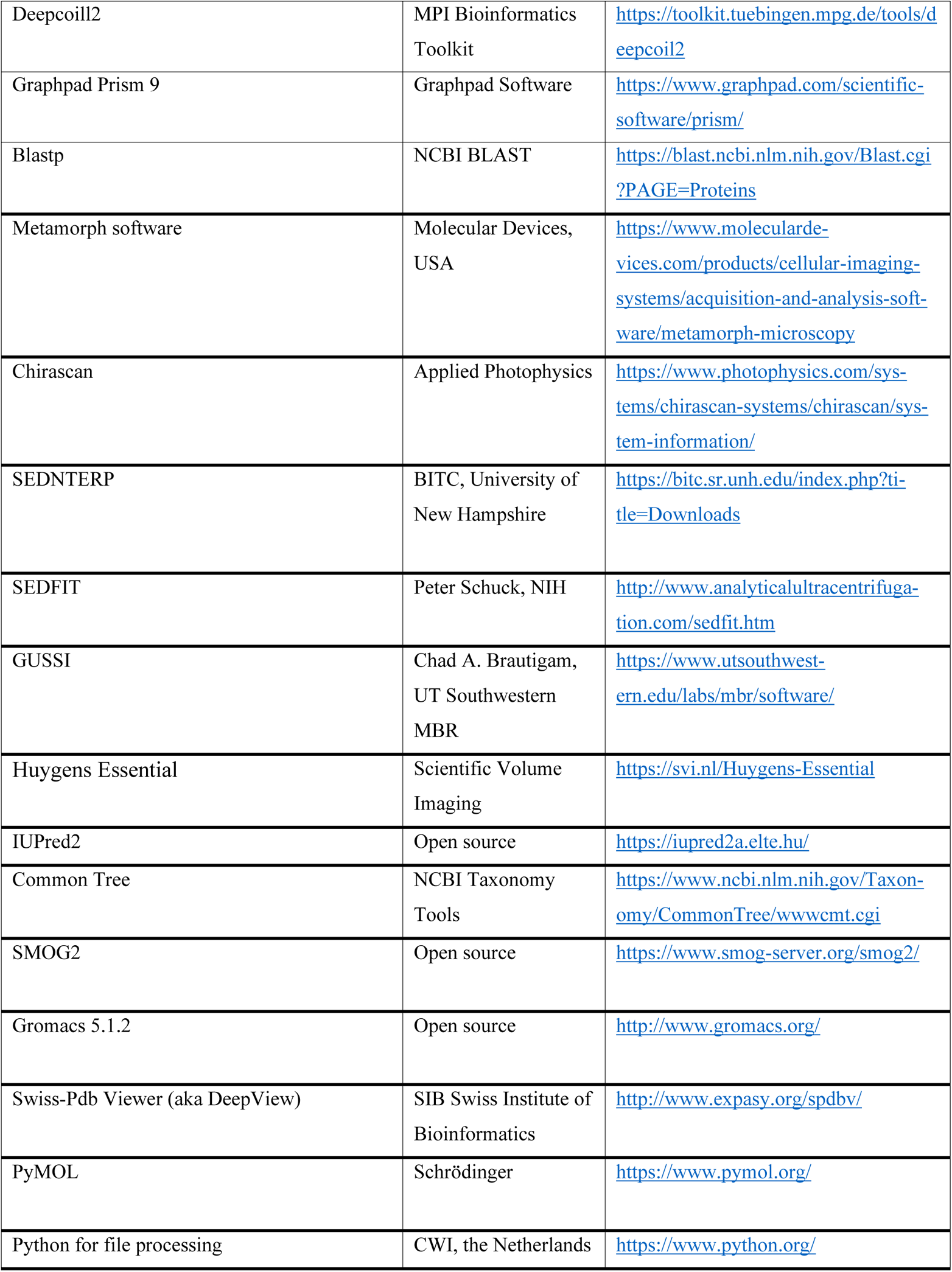
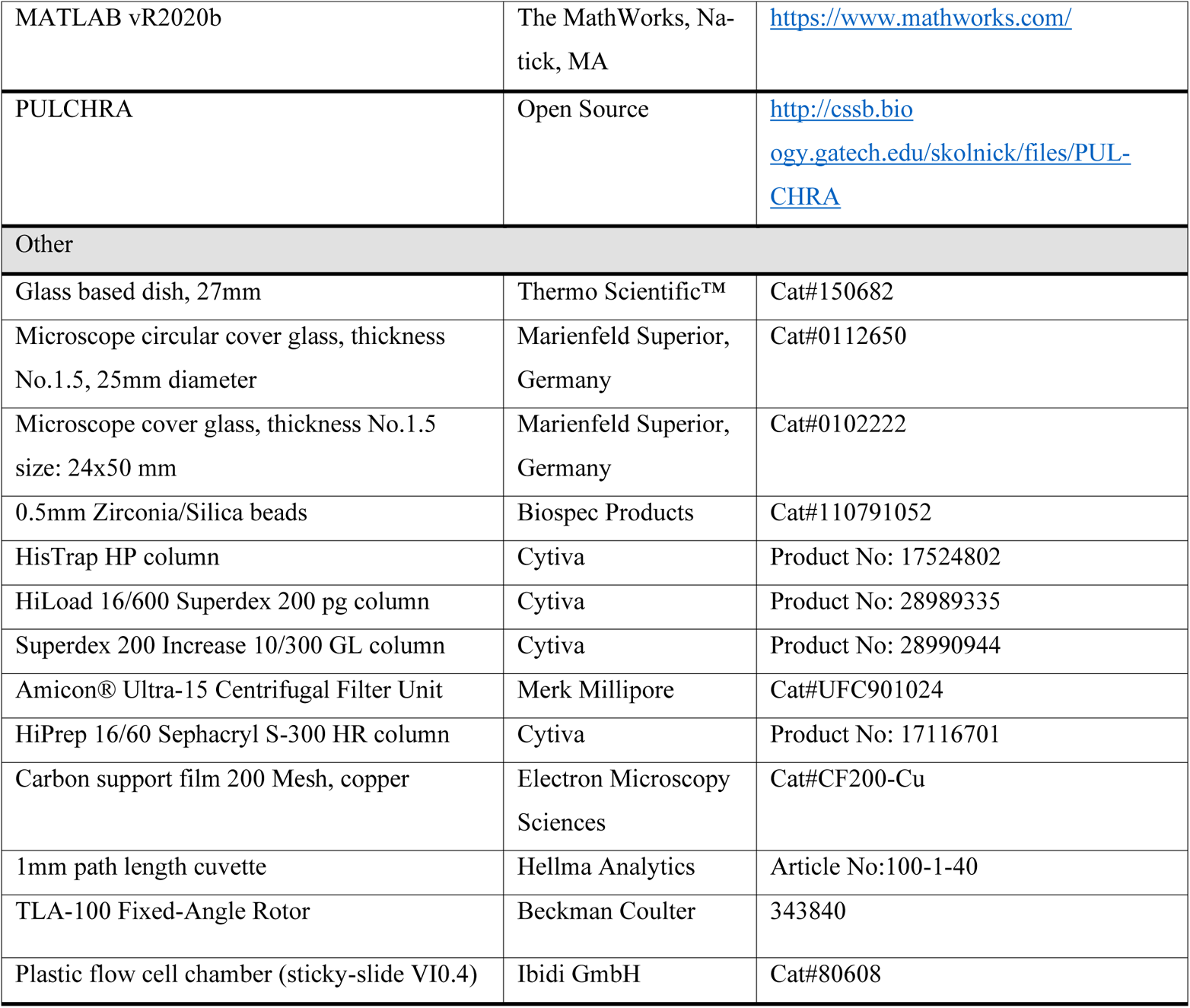

### RESOURCE AVAILABILITY

Further information and requests for resources and reagents should be directed to and will be fulfilled by Yansong Miao.

#### Materials availability

Yeast strains and plasmids created in this study are available upon request from the Lead Contact.

#### Data and code availability

This study did not generate any new large-scale data and code.

### EXPERIMENTAL MODEL AND SUBJECT DETAILS

#### Mouse embryonic fibroblast

Mouse embryonic fibroblast (MEF) was a gift from Dr. Satyajit Mayor from National Centre for Biological Sciences, Bangalore, India, and was used for lipofectamine transfection ^128^. Cells were grown in DMEM (high glucose, GlutaMAX™, Gibo) supplemented with 10% fetal bovine serum (GE hyclone), 100 units/ml penicillin and 100 µg/ml streptomycin at 37°C in CO_2_ incubator. To generate plasmids for overexpressing C-terminal GFP tagged Coronin1A and Coronin1C full-length (pEGFP-N1-Coro1A-FL and PEGFP-N1-Coro1C-FL) and truncation versions (pEGFP-N1-Coro1A-ΔIDR, pEGFP-N1-Coro1C-ΔIDR) in MEF cells, sequences encoding human Coronin1A and Coronin1C were amplified by PCR using cDNA clones from Mammalian Gene Collection (MGC) ^130^ as templates and inserted into pEGF-N1 vector. Plasmids of IDR truncation versions were generated by deleting 401aa-424aa and 401aa-433aa from full-length Coronin 1A and Coronin 1C plasmids, respectively. MEF cells were transiently transfected with plasmids above using Lipofectamine 3000 (Invitrogen, USA) following the manufacturer’s protocol and grown overnight on a round glass-bottom dish (Thermo Fisher) at 37°C in a CO_2_ incubator for protein expression. Transfected cells were identified by GFP fluorescence under microscope.

#### Saccharomyces cerevisiae strains

All yeast strains used in this study were derived from *S.cerevisiae* strain S288C ^131^ and listed in Table S1.

For spotting assay, strains were grown in YPD liquid medium (1% yeast extract, 2% peptone and 2% dextrose) at 25°C before plating on the YPD agar plate. For live-cell imaging, strains were grown in synthetic complete (SC) liquid medium with 2% dextrose without tryptophan at 30°C, or 25°C for *arc35-5* background strains, unless otherwise noted.

To construct plasmids for expressing mRuby2-tagged Crn1 full-length and truncation variants under native promoter in *S.cerevisiae*, pRS305-Crn1-FL-mRuby2-4*myc±500 plasmid was firstly generated by gene synthesis together with ClonExpress II (Vazyme Inc, China)-based cloning of sequences encoding full length Crn1, yeast-optimized mRuby2 ^132^ and 4xMyc, plus 500bp of upstream and downstream sequences of *CRN1,* which were inserted to integrating plasmid pRS305 in order ^133^, truncated variants (pRS305-Crn1-ΔN-mRuby2-4*myc±500, pRS305-Crn1-ΔCC-mRuby2-4*myc±500, pRS305-Crn1-ΔIDR-mRuby2-4*myc±500, pRS305-Crn1-N-mRuby2-4*myc±500, pRS305-Crn1-IDR-mRuby2-4*myc±500, pRS305-Crn1-CC-mRuby2-4*myc±500, and pRS305-Crn1ΔIDR-S-mRuby2-4*myc±500) were obtained by modifications on pRS305-Crn1-FL-mRuby2-4*myc±500 plasmid.

To express homo-oligomeric Crn1 variants, peptides of the dimer, trimer, tetramer, and pentamer coiled-coils ^61^ were synthesized from Biobasic (Bio Basic, Asia Pacific) and inserted at the original CC position of pRS305-Crn1-ΔCC-mRuby2-4*myc±500 plasmid. *Crn1Δ* strain YMY858 was obtained by replacing *CRN1* ORF with *Candida glabrata URA3* cassette in DDY904. The pRS305-derived plasmids were integrated into *LEU2* locus of strain YMY858 through XcmI digestion and lithium acetate transformation ^62^. Briefly, YMY858 was cultured in YPD medium at 30°C for overnight. The next day it was reinoculated into a new medium with a starting optical density (OD_600_)=0.2 and cultured until OD=0.8-1. 10 mL cultures were harvested by centrifugation for 10 min at 2000 x *g* at room temperature and washed once with sterile, double-distilled water. Yeasts were resuspended in 0.1 ml yeast transformation resuspension buffer (100 mM lithium acetate, 10 mM Tris, pH 8.0, 1 mM EDTA, pH 8.0) with 2-5µg plasmids pre-digested by XcmI and 50 µg of herring sperm carrier DNA (Promega), then 700 µL of yeast transformation buffer (40% (w/v) polyethylene glycol, 100 mM lithium acetate, 10 mM Tris, pH 8.0, 1 mM EDTA, pH 8.0) was added and mixed well, the mixture was incubated at 30°C for 30 min followed by heat shock in 42°C water bath for additional 30 min. The mixture was centrifuged again for 5 min at 2000 x *g* to collect yeast cells, and cells were then resuspended with 200 µL sterile, double-distilled water and spread on a synthetic complete (SC) agar plate with 2% dextrose without leucine to select positive transformants. To introduce various *CRN1* variants into the *arc35-5* background, we crossed them with strain YXD160 (*crn1Δ arc35-5)*, diploids were picked and induced to sporulate, tetrads were dissected and plated on YPD agar plate with 200 μg/ml Geneticin and SC ager plate with 2% dextrose without leucine to select target strains.

#### Yeast live-cell imaging

Yeast strains were cultured in the SC liquid medium with 2% dextrose without tryptophan overnight and re-inoculated into a new medium to a staring OD_600_=0.2. Cells were allowed to grow for additional 4 hours before imaging. Cells were immobilized on concanavalin A (1mg/ml, Sigma)-coated circular coverslip (Marienfeld Superior) and imaged at 25°C by a wide-field microscope Leica Dmi8 (Leica Microsystems) equipped with ORCA-Flash 4.0 LT scientific CMOS camera (Hamamatsu Photonics, Japan) and Leica ×100 oil immersion objective lens (NA 1.4) using Metamorph software (Molecular Devices). Images were acquired as a z-axis stack with a step size of 0.25µm for a total of 31 frames. Middle focal panel images were used for representatives and intensity analysis.

#### Yeast growth assay

For the yeast spotting assay, each strain was inoculated into YPD liquid medium and cultured at 25°C for overnight. The next day, the statured culture was inoculated to a fresh YPD medium with a starting OD_600_=0.2 and cultured for additional 3 hours. An additional reinoculation was performed starting from OD_600_=0.2 and cultured for another 3 hours before being diluted to OD_600_=0.1 for spotting assay with ten-fold serial dilutions in YPD medium. 4 μL culture of each dilution was spotted on the YPD agar plate. Plates were incubated at the tested temperatures for 48 hours and scanned by a Perfection V600 Photo scanner (Epson) with 600 dpi. Images were converted to grayscale and inverted in Photoshop.

#### Yeast whole-cell extraction and immunoblotting

Yeast whole-cell protein extraction was prepared by trichloroacetic acid (TCA) precipitation. Yeasts were cultured in YPD liquid medium overnight at 30°C or 25°C for *arc35-5* background strains. Cells were reinoculated to OD_600_=0.2 and cultured until OD_600_=0.8-1. 10 OD cells were collected by centrifugation at 2000 x *g* for 10 min and washed once with sterile, double-distilled water. Cells were resuspended in 250 µL of 20% TCA (Sigma) solution with 100 µL 0.5mm Zirconia/Silica beads (Biospec Products) and lysed by Powerlyzer homogenizer (Qiagen) at 3500 rpm for 1 min, which was repeated 3 times with 5 min interval on ice. Lysates were transferred out, and beads were washed once with 300 µL of 5% TCA to collect the remaining lysates. An additional 700 µL of 5% TCA solution was added to lysates to reach final around 8% TCA. Samples were centrifuged at 14,000 x *g* for 10 min in an Eppendorf™ Benchtop Microcentrifuge (Thermo Fisher). Pelleted proteins were resuspended in 40 µL of 1M Tris (pH 8.0) and 80 µL of 2X sodium dodecyl sulfate (SDS) loading buffer (100 mM Tris, pH 6.8, 5% SDS, 20% glycerol, 0.1% bromophenol blue, and 200 mM DTT) and boiled at 95°C for 10 min. Samples were centrifuged again at 14,000 x *g* for 10 min, and supernatants were collected for SDS-PAGE and western blot detecting. Primary antibodies used for western blot: monoclonal mouse anti-pgk1(22C5D8, Invitrogen) was diluted in TBST (20 mM Tris, pH 7.5, 150 mM NaCl, 0.1% v/v Tween20) as 1:1000, monoclonal mouse anti c-Myc (9E10, self-prepared) was diluted in TBST as 1:1000. Secondary antibodies: IRDye® 800CW Goat anti-Mouse (LI-COR) was diluted in TBST as 1:10,000. Blots were scanned by an Odyssey Infrared Imager (LI-COR Biosciences).

#### Mammalian cell imaging

To image subcellular localization of GFP-tagged Coronin1A or Coronin1C protein variants and filamentous F-actin in MEF cells, cells were washed twice with prewarmed phosphate-buffered saline buffer (PBS, pH 7.4), and fixed with 4% paraformaldehyde (PFA) solution in PBS for 20 min at room temperature, and fixed cells were permeabilized with 0.1% Triton X-100 (Bio-Rad) in PBS for 5min. After washing twice with PBS, Alexa Fluor™ 647 Phalloidin (Invitrogen) (1:40 dilution in PBS with 1% bovine serum albumin) was applied to cells for 20 min at room temperature in the dark area, and cells were washed three times with PBS. Samples were imaged by a wide-field microscope Leica DMi8 (Leica Microsystems) equipped with ORCA-Flash 4.0 LT scientific CMOS camera (Hamamatsu Photonics, Japan) with Leica ×100 oil immersion objective lens (NA 1.4) controlled by Metamorph software (Molecular Devices). Z-axis scanning images were acquired with a step size of 0.25 µm for a total of 31 frames. An image of the middle focal plane was shown as the representative image. Images were deconvolved by Huygens Essential deconvolution software (Scientific Volume Imaging, Netherlands).

#### Protein expression and purification

A series of yeast Crn1 protein expression plasmids were constructed for protein expression and purification from *E.coli* Rosetta (DE3) using a pET-21d(+)(pSY5) vector that contains an 8x histidine tag and an HRV 3C protease cleavage site at the N-terminus of the protein of interest ^129^. To generate pSY5-Crn1-FL, pSY5-Crn1-N, pSY5-Crn1-IDR, pSY5-Crn1-ΔN, and pSY5-Crn1-ΔCC plasmids, Crn1 1-651aa, 1-400aa, 401-604aa, 401-651aa, and 1-604aa were amplified by PCR, respectively, and inserted to pSY5 backbone through T4 ligation. pSY5-Crn1-ΔIDR plasmid keeping 401aa-604aa was generated from pSY5-Crn1-FL plasmid, whereas pSY5-Crn1-CC plasmid was generated by truncating 401-604aa from the pSY5-Crn1-ΔN plasmid. The pSY5-Crn1-ΔIDR-S plasmid was generated by a two-step truncation that excluded 401aa-483aa and 524aa-601aa from the pSY5-Crn1-FL plasmid. Homo-oligomeric Crn1 constructs were obtained by inserting dimer, trimer, tetramer, and pentamer peptides ^61^ at the original CC position of the pSY5-Crn1-ΔCC plasmid. pSY5-Las17-polypVCA plasmid was constructed to express 300-633aa of Las17 ^65^.

pSY5-based plasmids were transformed to *Escherichia coli* Rosetta (DE3) through 1.5 min heat shock in 42°C water bath, bacteria were then spread on LB agar plate (1% tryptone, 0.5% yeast extract, 1% NaCl, and 1% agar) with 100 µg/ml ampicillin to select positive colonies. A single colony was picked to 20 mL LB liquid medium and grown at 37°C for overnight. Culture was scaled to 1 L in TB liquid medium (2.4% yeast extract, 2% tryptone, 0.4% glycerol, 0.017M KH_2_PO_4_ and 0.072M K_2_HPO_4_) with 100 µg/ml ampicillin and 35 µg/ml chloramphenicol. When OD_600_ reached 1.5, expression was induced by 0.5 mM IPTG at 16°C for overnight. Cells were harvested by centrifugation at 4°C, 5000 x *g* (rotor JA10, Beckman Coulter) and lysed by LM20 microfluidizer (20000 psi) in binding buffer (50 mM Tris, pH 8.0, 500 mM NaCl, 20 mM Imidazole) with 1mM PMSF and protease inhibitor tablet (Thermo Fisher). Lysates were centrifuged at 20,000 x *g*, 4°C for 1 hour (JA 25.5 rotor, Beckman Coulter). The supernatant was kept and clarified by a 0.45 µM syringe filter (Pall Corporation), then loaded onto a 5 mL Histrap column (Cytiva) connected to an FPLC AKTAxpress system (GE Healthcare). After affinity binding, the column was washed with binding buffer followed by gradient elution from 20 mM to 500 mM Imidazole using elution buffer (50 mM Tris, pH 8.0, 500 mM NaCl, 500 mM imidazole). Peak fractions were checked by SDS-PAGE, good fractions were combined, and dialyzed in protein buffer (50 mM Tris, pH 8.0, 150 mM NaCl) overnight at 4°C. When necessary, the dialyzed protein solution was further separated by size exclusion chromatography using a HiLoad 16/600 superdex 200 pg column (Cytiva) pre-equilibrated with protein buffer. Peak fractions were collected and checked by SDS-PAGE, then good fractions were combined and concentrated with 10-kDa or 30-kDa cut-off concentrators (Merk Millipore). Protein concentration was measured by the BCA protein assay kit (Thermo Fisher). The protein solution was aliquoted into a small volume, frozen in liquid N2 and stored in −80°C freezer for storage.

#### Rabbit skeletal muscle actin (RMA) purification and labeling

To obtain monomeric ATP-bound RMA for actin cosedimentation assay, transmission electron microscopy, and TIRF microscopy. 2g rabbit muscle acetone powder (Pel-Freez, LLC) was dissolved in a 200 mL cold G-buffer (2 mM Tris, pH 8.0, 0.2 mM ATP, 0.5 mM DTT and 0.1 mM CaCl_2_) and stirred at 4°C overnight. The mixture was filtered with a cheese cloth to remove muscle powder, then the actin-dissolved solution was further centrifuged at 2600 x *g*, 4°C (Type 45 Ti rotor, Beckman Coulter) for 30 min to collect supernatant. Actin in supernatant was then polymerized with slowly stirring for 1 hour at 4°C by adding KCl and MgCl_2_ solution to a final concentration of 50 mM and 2 mM, respectively. To remove tropomyosin and other actin binding proteins, fine KCl powder was slowly added to reach a final concentration of 0.6 M and the solution was kept stirring for another 30 min. The solution was then centrifuged at 14,000 x *g* (Type 45 Ti rotor, Beckman Coulter) for 3 hours at 4°C to collect the filamentous actin pellet. Pellet was then rinsed with cold G-buffer and homogenized with a homogenizer in 7 mL of cold G-buffer followed by a short time of sonication. The sample was then dialyzed in 2 L G-buffer at 4°C for 48 hours to induce depolymerization (G-buffer was changed every 12 hours). After buffer exchange, the sample was centrifuged at 167,000 x *g* (SW 55 Ti swinging-bucket rotor, Beckman Coulter) at 4°C for 2.5 hours, 5 mL of supernatant was collected and loaded to a Sephacryl S-300 HR column (GE healthcare) pre-balanced with G-buffer. Peak fractions were collected and combined, then 0.01% (final) sodium azide (Sigma) was added to inhibit fungi contamination and kept at 4°C. Actin concentration was measured by reading OD_290_ with a Nanodrop 2000 (Thermo Scientific).

To label actin with Oregon Green™ 488 Iodoacetamide (Invitrogen) or NHS-dPEG®4-biotin (Sigma), the same steps were followed as RMA purification until pelleted filamentous actin was homogenized and sonicated. After this, the sample was dialyzed in a 1 L G-buffer at 4°C for overnight. The next day, the sample was changed to 1 L G-buffer without DTT and dialyzed for 4 hours at 4°C (buffer changed once). Oregon Green™ 488 Iodoacetamide or NHS-dPEG®4-biotin was dissolved in dimethylformamide to a final concentration of 10 mM. Before labeling, actin concentration was measured by reading OD_290_ with a Nanodrop 2000 (Thermo Scientific). Actin was firstly diluted with an equal volume of cold 2X labeling buffer (50 mM Imidazole, pH 7.5, 200 mM KCl, 0.6 mM ATP and 4 mM MgCl_2_) and further diluted to final 23 µM with cold 1X labeling buffer, then 10-fold molar excess of Oregon Green™ 488 Iodoacetamide or NHS-dPEG®4-biotin was added dropwise while very gently vortexing. The mixture was covered with aluminum foil and rotated at 4°C for overnight. The next morning, labeled filamentous actin was centrifuged at 167,000 x *g* (Type 50.2 rotor, Beckman Coulter) for 3 hours at 4°C. Pellets were collected and homogenized in 4 mL G-buffer, waited on ice for 1 hour, and homogenized again. Actin was then dialyzed in 1 L G-buffer at 4°C for 48 hours to induce depolymerization (dark, G-buffer changed every 12 hours). After buffer exchange, actin was centrifuged at 436,000 x *g* (TLA100 rotor, Beckman Coulter) at 4°C for 1 hour. The supernatant was collected and further purified by Sephacryl S-300 HR column (GE healthcare) pre-balanced with G-buffer. Peak fractions were collected and combined, then dialyzed in a 500 mL G-buffer with 50% (v/v) glycerol at 4°C overnight to reduce volume. Small aliquots were frozen in liquid nitrogen and stored in a −80°C freezer.

#### Yeast protein expression and purification

To generate plasmid for overexpression C-terminal 6xHis-tagged yCrn1-FL protein in budding yeasts, DNA sequences encoding Crn1 full length was amplified using pRS305-Crn1-FL-mRuby2-4*myc±500 as a template and inserted into the pYeast Pro vector (pGAL-ORF-3×StreptagII-9×His)^112^. Plasmids were transformed into strain YMY2043^112^ using lithium acetate, and yeasts were spread on an SC agar plate with 2% dextrose without uracil to select positive transformants.

To obtain large-scale yeast culture for protein purification, we followed a published protocol^62^, 10-20 positive transformants were inoculated into 10 mL SC medium with 2% raffinose without leucine and grow at 30°C with vigorous shaking for around 2 days until saturation (OD_600_ 2 to 3). Then statured culture was scaled to 100 mL using SC medium with 2% raffinose (MP biomedicals) without leucine and grew at 30°C with vigorous shaking until saturation again. The culture was then transferred to 1.9 L of fresh SC medium with 2% raffinose without leucine in a 5-L flask and kept growing at 30°C. When the culture reached saturation after 36 to 48 hours, 160 mL of 30% (w/v) galactose (Sigma) and 240 mL of 10X YP (10% w/v yeast extract, 20% w/v peptone) were added to the flask making the final concentration of 2% galactose and 1X YP, in which protein was induced to express for 12 to 16 hours with vigorous shaking at 30°C. Yeasts were collected by centrifugation at 6,000 x *g* (rotor JA10, Beckman Coulter) for 15 min at 4°C, and washed twice with sterile double-distilled water. Cells were then resuspended with 20% volume of sterile, double-distilled water, the mixture was frozen into small balls by dripping into liquid nitrogen, which was further thoroughly ground by a Freezer/Mill 6870 cryomilling machine (SPEX SamplePrep) into a fine powder and stored at −80 °C. For protein purification, 5 grams of yeast powder was dissolved in 30 mL binding buffer with 50 µL protease inhibitor cocktail IV (BioWorld), and 300 µL of 100 mM PMSF, 75 µL of 0.2 M sodium orthovanadate (Sigma), 300 µL of 1 M glycerophosphate (Sigma) and 600 µL of 0.5 M NaF. The mixture was centrifuged at 20,000 x *g* (JA 25.5 rotor, Beckman Coulter) at 4 °C for 1 hour. The supernatant was isolated and clarified by a 0.45 µm syringe filter (Pall Corporation). Filtered supernatant was loaded into a 5 mL Histrap column (Cytiva) connected to an FPLC AKTAxpress system (GE Healthcare). After affinity binding, the column was washed with binding buffer followed by gradient elution from 20 mM to 500 mM Imidazole using elution buffer. Peak fractions were collected and checked by SDS-PAGE, fractions were combined and dialyzed in 2 L protein buffer (50 mM Tris, pH 8.0, 150 mM NaCl) at 4°C for overnight, then concentrated with 50-kDa cut-off concentrators (Merk Millipore). Protein concentration was measured by the BCA protein assay kit (ThermoFisher). For storage, the protein was aliquoted into a small volume, frozen in liquid N_2,_ and stored in −80°C freezer.

#### Specimen preparation and transmission electron microscopy

Purified RMA was polymerized for 1 hour in F-buffer (2 mM Tris, pH 8.0, 50 mM KCl, 1 mM MgCl_2_, 1 mM EGTA, 0.2 mM ATP, 0.5 mM DTT, and 0.1 mM CaCl_2_). 1 µM of each purified different Crn1 protein variant was incubated with 2 µM of pre-polymerized F-actin for 30 min. Samples were then applied to glow discharged carbon-coated copper grid (200 mess, Electron Microscopy Sciences) and waited for 2 min. Extra volume was removed by filter paper, and grids were negatively stained with 1% uranyl acetate (Electron Microscopy Sciences) for 1 min. Grids were air-dried and examined at 120kV by an FEI Tecnai 12 TEM equipped with an Ultrascan 1000 CCD camera (Gatan, Inc).

#### Far-UV circular dichroism

Purified recombinant Crn1-CC and Crn1-IDR proteins were dialyzed against 50 mM sodium phosphate buffer (pH 7.4, 37.7 mM Na_2_HPO_4,_ 12.3 mM NaH_2_PO_4_) at 4°C for overnight. Protein was diluted to a final concentration within the optimal range for the detector. 300 µL of the sample was loaded into a 1 mm path length quartz cuvette (Hellma Analytics), spectra were recorded on a Chirascan^TM^ Circular Dichroism Spectrometer (Applied Photophysics) equipped with a temperature controller at 20°C and supplied with constant N_2_ flushing. Spectra were acquired from 260 nm to 190 nm (step size: 1 nm) with an integration time of 1sec at each wavelength, and baseline was corrected by a sample with buffer alone. Raw data in machine units θ (mdeg) were converted to Mean Residue Ellipticity [θ] (degrees cm^2^ dmol^-1^ residue^-1^) by equation: [θ] = θ x (0.1 x MRW)/(P x conc), where MRW=protein weight (daltons) / number of residues, P=patch length (cm) and conc=protein concentration (mg/ml).

#### Actin cosedimentation assays

For high-speed actin cosedimentation assay, a range of concentrations of Crn1 protein variants was incubated in F-buffer with pre-polymerized actin filaments at room temperature for 30 min and then spun at 100,000 *× g* (high speed) with TLA100 rotor (Beckman Coulter) for 20 min at 25°C. For low-speed actin cosedimentation assay, a range of concentrations of Crn1 protein variants was incubated in a buffer with physiological ionic strength (20 mM Tris, pH 8.0, 150 mM KCl, 1 mM MgCl_2_, 1 mM EGTA, 0.2 mM ATP, 0.5 mM DTT, and 0.1 mM CaCl_2_) with pre-polymerized actin filaments at room temperature for 30 min, and then spun at 10,000 *× g* (low speed) using an Eppendorf™ Benchtop Microcentrifuge (Thermo Fisher) at 25°C. For both assays, an equal volume of supernatant and pellet fractions were collected and analyzed by SDS-PAGE and stained by GelCode™ Blue Stain Reagent (Thermo Fisher) overnight. Gels were imaged by a Perfection V600 Photo scanner (Epson), and densitometry of the band was quantified by ImageJ. For the high-speed assay, a total (before centrifugation) fraction was also collected for SDS-PAGE for making a standard curve.

#### Total internal reflection fluorescence microscopy

24×50-mm glass coverslips (Marienfeld Superior) were cleaned by immersion in 20% sulfuric acid (Sigma) for overnight and rinsed thoroughly with double-distilled water 3 times. Cleaned coverslips were then coated with final 2 mg/ml methoxy-PEG-silane and 2 µg/ml biotin-PEG-saline (Laysan Bio Inc) in 80% ethanol (pH 2.0 with HCl) in 70°C water bath for overnight. The next day, coverslips were rinsed thoroughly with double-distilled water and dried with an N_2_ stream. Coated coverslips were covered with aluminum foil and stored in −80°C. Before experiments, a coated coverslip was stuck to the bottom side of a plastic flow cell chamber (Ibidi, sticky-slide VI0.4). Then 30 µL of HBSA (20 mM HEPES, pH 7.5, 1 mM EDTA, 50 mM KCl, and 1% bovine serum albumin) was pipetted to a cell and incubated for 1 min, then 30 µL of 0.1 mg/ml streptavidin in HEKG100 (20 mM HEPES, pH 7.5, 1 mM EDTA, 50 mM KCl, and 10% (v/v) glycerol) was pipetted and waited for additional 1 min. Afterwards, 30 µL solution was pipetted out, cell was washed by 30 µL of 1X TIRF buffer (10 mM Imidazole, pH 7.4, 50 mM KCl, 1 mM MgCl2, 1 mM EGTA, 0.3 mM ATP, 50 mM DTT, 15 mM glucose, 400 μg/ml glucose oxidase, 40 µg/ml catalase, and 0.5% methylcellulose [4000 cP]) for two times. Proteins were mixed with RMA (20% Oregon Green 488 labeled, 1% biotin-labeled) in G-buffer to designated concentration, and 30 µL was pipetted to cell followed by 30 µL of 2X TIRF buffer to initiate polymerization. To image Crn1-bundled actin filaments, a high salt 1X TIRF buffer with physiological ionic strength (10 mM imidazole, pH 7.4, 150 mM KCl, 1 mM MgCl2, 1 mM EGTA, 0.3 mM ATP, 50 mM DTT, 15 mM glucose, 400 μg/ml glucose oxidase, 40 µg/ml catalase, and 0.5% methylcellulose [4000 cP]) was used. Images were acquired as a stack at room temperature with 30-sec intervals for 15 min (for Arp2/3-mediated actin polymerization assay) or 5 min interval for 90 min (for Crn1-mediated actin-bundling assay) using a Nikon Ti2-E inverted microscope equipped with a 100x 1.45NA Plan-Apo objective lens and a TIRF module (iLasV2 Ring TIRF, GATACA Systems) and an ORCA-Fusion sCMOS camera (Hamamatsu Photonics). Imaging lasers were provided by 405nm/100mW (Vortran), 488nm /150mW (Vortran), 561nm /100mW (Coherent) and 639nm /150mW (Vortran) combined in a laser launch (iLaunch, GATACA Systems). Focus was maintained by hardware autofocus (Perfect Focus System), and image acquisition was controlled by MetaMorph software (Molecular Device). For Arp2/3-mediated actin polymerization assay, images were deconvolved by Huygens Essential deconvolution software (Scientific Volume Imaging, Netherlands) to diminish background signal.

#### Analytical ultracentrifugation (AUC)

AUC Sedimentation Velocity (AUC-SV) experiments were performed on Beckman Proteome Lab XL-I Analytical Ultracentrifuge using an 8-hole An-50 Ti analytical rotor. Samples were dialyzed overnight in physiological buffer (50 mM Tris, pH 8.0, 150 mM NaCl) and loaded into 2-sector cells fitted with 1.2 cm epon centerpiece and quartz windows. The samples were centrifuged at 30,000 rpm or 45,000 rpm at 20 °C and absorbance at 280 nm (for Crn1-FL, Crn1-ΔCC and yCrn1-FL) and 230 nm (for Crn1-CC, Crn1-IDR, and Crn1-ΔN) was recorded every 5-10 minutes during 15 hours centrifugation. The data were analyzed with SEDFIT using *c(s)* and *c(s,ff_0_)* size distribution models ^134^, and plotted with GUSSI ^135^. Sedimentation coefficients were standardized to s_20,w_ using the partial specific volume of the proteins (calculated using SEDFIT), solvent density, and viscosity (calculated using SEDNTERP) ^136^.

#### Coronin sequence analysis and the taxonomic tree

The murine Coronin1A (NCBI accession NP_034028.1) N-terminal sequence (1-402aa) was input as a query sequence to NCBI blastp against non-redundant protein sequence database to identify coronin homologues for the 200 species that contain coronin family members, which were previously reported in ^66^.

We firstly excluded the “hypothetical/unnamed/predicted/uncharacterized/putative” candidates. In the remaining list of coronin candidates, we defined a cut-off for “total alignment score” of 350, allowing us to identify the coronin homologues containing around N-terminal 400aa, which is the core and conserved coronin N-terminal domain. Thereby, we have identified 392 coronin homologues from 200 species. For each identified coronin homologue, DeepCoil2 (https://toolkit.tuebingen.mpg.de/tools/deepcoil2) ^54^ was used to predict coiled coil (CC) region, from which 11 (one from *B.bigemina,* one from *P.berghei*, one from *P.chabaudi,* one from *P.knowlesi,* one from *P.reichenowi,* one from *P.Vivax,* one from *P.yoelii,* one from *T.annulate*, one from *A.aegypti,* and two from *M.lucifugus*) out of 392 coronin homologues do not have predicted CC. The unique regions (IDR) were identified between N-terminal and CC which were used to generate IDR-MAP. To build the taxonomic tree of the 200 species, taxid of each species was input to NCBI taxonomy tool-common tree, and generated tree was modified by iTOL (https://itol.embl.de/) ^126^. The length of the predicted IDR was then integrated as a bar diagram outside the taxonomy tree, resulting in an IDR-MAP.

#### Coarse-grained Simulation

##### Coarse-grained (CG) model

The CG model represents every amino acid residue with one bead at its Cα atom position, following Ravikumar et al.^137^, originating from the structure-based (i.e., Gō-like) coarse-grained force field ^138^. The potential energy function includes the interactions for bonds (E_bond_), angles (E_angle_), dihedral angles (E_dihedral_), native-like contacts modeled by Lennard-Jones (LJ)-type potentials (E_native_), electrostatic terms (E_elec_) and hydrophobic terms (E_H_) for the structured part of protein, namely,

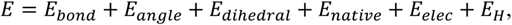

where 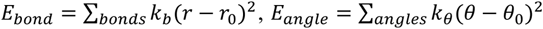 and 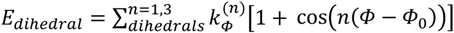. Intrinsically disordered region (IDR, defined according to the sequence analysis above) neglects the angular and dihedral terms for flexibility. Here, r, θ and Φ are instantaneous bond lengths, angles and dihedral angles, respectively, while *r*_0_, *θ*_0_, and *ϕ*_0_ are the corresponding values in the reference crystal structure. Force constants *k_b_* = 4.184 × 10^4^ kJ/mol, *k_θ_* = 8.368 × 10 *kJ*/mol · *rad*, 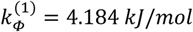, and 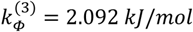.

The LJ-type potential energy function for native contacts is used between residue i and residue j as

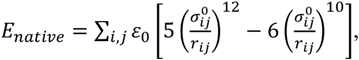

where *r_ij_* is the distance between residue *i* and *j*, while 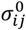 is the corresponding distance in the reference crystal structure. The native contact pairs are generated by the software SMOG2 based on calculations using the atomistic structure ^139^.

The electrostatic (E_elec_) term is denoted as

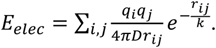

Here, the electrostatic potential energy is treated through the Debye-Hückel method ^140^. *q_i_* represents the charge of residue *i*. The Debye screening length *k* equals 1 nm, corresponding to an ionic strength of approximately 100 mM, and the dielectric constant D equals 80 for the solvent medium (water). At pH 7.0, residue charges *q_i_* = +1*e* for Lys and Arg, *q_i_* = −1*e* for Asp and Glu, and *q_i_* = +0.5*e* for His (*e* represents the elementary charge).

The hydrophobic interactions (E_H_) are either attractive (*ε_ij_* < 0) or repulsive (*ε_ij_* ≥ 0), where

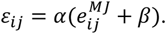

Here, 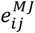 is the Miyazawa-Jernigan (MJ) contact energy between residue *i* and residue *j* ^141^, α is the parameter to scale E_H_ in relation to E_elec_, and β (in the unit of RT, where R is the ideal gas constant and T represents temperature in Kelvin) is the parameter to balance the attractive and repulsive interactions ^137^. Therefore, the hydrophobic interactions 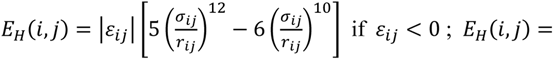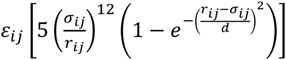 where *d* is equal to 0.38 nm. Finally, a scaling factor γ is introduced for *σ_ij_* as

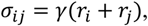

where *r_i_* is the van der Waals radius of residue *i* ^142^.

##### Langevin Dynamics (LD) Simulation

The CG model was implemented for Langevin dynamics (LD) simulations with the GROMACS 5.1.2 package ^143^. All simulations were carried out in the NVT ensemble (constant atom number, simulation box size, and temperature). Simulations were performed at 300K with a friction coefficient of 50/ps. A simulation time step of 0.01ps was used. At least 1000 ns LD runs were conducted. Structures for Crn1 coiled-coil and IDRs were modeled by the software spdbv (http://www.expasy.org/spdbv/). The structure for Coro1A coiled-coil was from Protein Data Bank (PDB ID: 2akf). The initial structure for each simulation was constructed with PyMOL ^144^.

##### Data analysis

The potential energy per chain was used for evaluating the stability of different oligomers, and energy contributions from electrostatic and hydrophobic interactions were further analyzed. For a certain kind of species, e.g., murine Coronin1A coiled-coil (MmCC), its monomer was taken as the reference state whose potential energy per chain was shifted to zero.

All the contact maps followed the workflow demonstrated in Figure 7-figure supplement 3. The motif-averaged contact map (Figure 7-figure supplement 2D, G) was created to present the residue-residue contact pattern. Each motif box, a 7×7 matrix (marked as a red square in Figure 7-figure supplement 3A) indicates the contact between the regions where each wholesome motif locates at two chains. Each wholesome motif was predicted from the sequence according to the coiled-coil packing pattern. Each motif-averaged contact map, a 7×7 matrix also, is the mean matrix for all the motif boxes. The motif-averaged contact map represents the average contact probability between different pairs of motif sites. As none of the contact probability values are larger than 0.5, the maximum value on the scale of contact probability is changed from 0 to 0.5 for better contrast. Following the approach of Ryan et al. ^145^, the inter-residue distance cut-off for a contact to be formed is *σ_ij_* as defined above. In addition to the original motif-averaged probability map, a motif contact difference map (Figure 7-figure supplement 2E, F; Figure 7I and J) was constructed to present the contact probability differences at different contact pairs for two proteins (e.g. ScIDR-MmCC and MmCC in Figure 7-figure supplement 3B). The motif contact difference map indicates the difference in each contact pair between protein A and protein B, denoted as “motif contact difference map (A minus B)”. Finally, the Euclidean distance was calculated as the score to measure the difference between two contact probability maps.

All the analysis was based on the last 500 ns of each simulation. GROMACS tools and MATLAB (v. R2020b; The MathWorks, Natick, MA) were used to analyze simulation trajectories. Protein 3D structures were rendered by PyMOL.

### QUANTIFICATION AND STATISTICAL ANALYSIS

#### mRuby2-patch intensity analysis

As illustrated in Figure 3-figure supplement 1A, to measure mRuby2 signal intensity at the actin patch, a rectangular box at 36×8 pixels was drawn crossing the Crn1-localized patches. The box was placed perpendicularly to the mother cell membrane, with a patch in the middle. Cytosol region was defined by proximal 8×8 box within the cell, patch region was defined by 8×8 box in the middle, and background region was defined by distal 8×8 box outside of the cell. The average intensity of each box was measured as signal intensity, termed as cytosol intensity (C), patch intensity (P), and background intensity (B), separately. Patch to total ratio was calculated use the equation: (P-B)/ [(P-B) +(C-B)].

#### Actin cosedimentation analysis

The SDS-PAGE gel was scanned by the scanner and converted to an 8-bit gray value image in ImageJ. A rectangle area covering the entire band was selected, area size and a mean gray value (mgv) were obtained from ImageJ, whereas a same-sized rectangle box from the background area was also selected for normalization. Band density was calculated as area x [(255-sample mgv) - (255-background mgv)]. F-actin bound Crn1 protein variants in high-speed actin cosedimentation assay were analyzed as previously described ^146^. To calibrate measured signal intensity by protein quantity, we firstly generated a standard curve by plotting the band density versus protein mass after measuring the gel of the total fraction. The measured band density of F-actin bound Crn1 from pellet fraction was converted to Crn1 amount (mass) using the standard curve after subtracting the Crn1 band density in the control experiments without F-actin. Data points were curve fitted using the Hill equation in GraphPad Prism 9. To analyze actin bundling in low-speed cosedimentation assay, both pellet and supernatant fractions of actin were used for analysis: at each Crn1 concentration, actin band densities from the pellet (P) and supernatant (S) were measured, percentage of actin in the pellet (%) was calculated as P/(P+S).

#### Statistical Analysis

Statistical analyses were performed in GraphPad Prism 9, unpaired two-tailed Student’s t test assuming equal variances was used to determine difference between 2 groups, *p <0.05; **p < 0.01; ***p < 0.001; ****p < 0.0001, ns = not significant.

---

**Figure 1–figure supplement 1.**
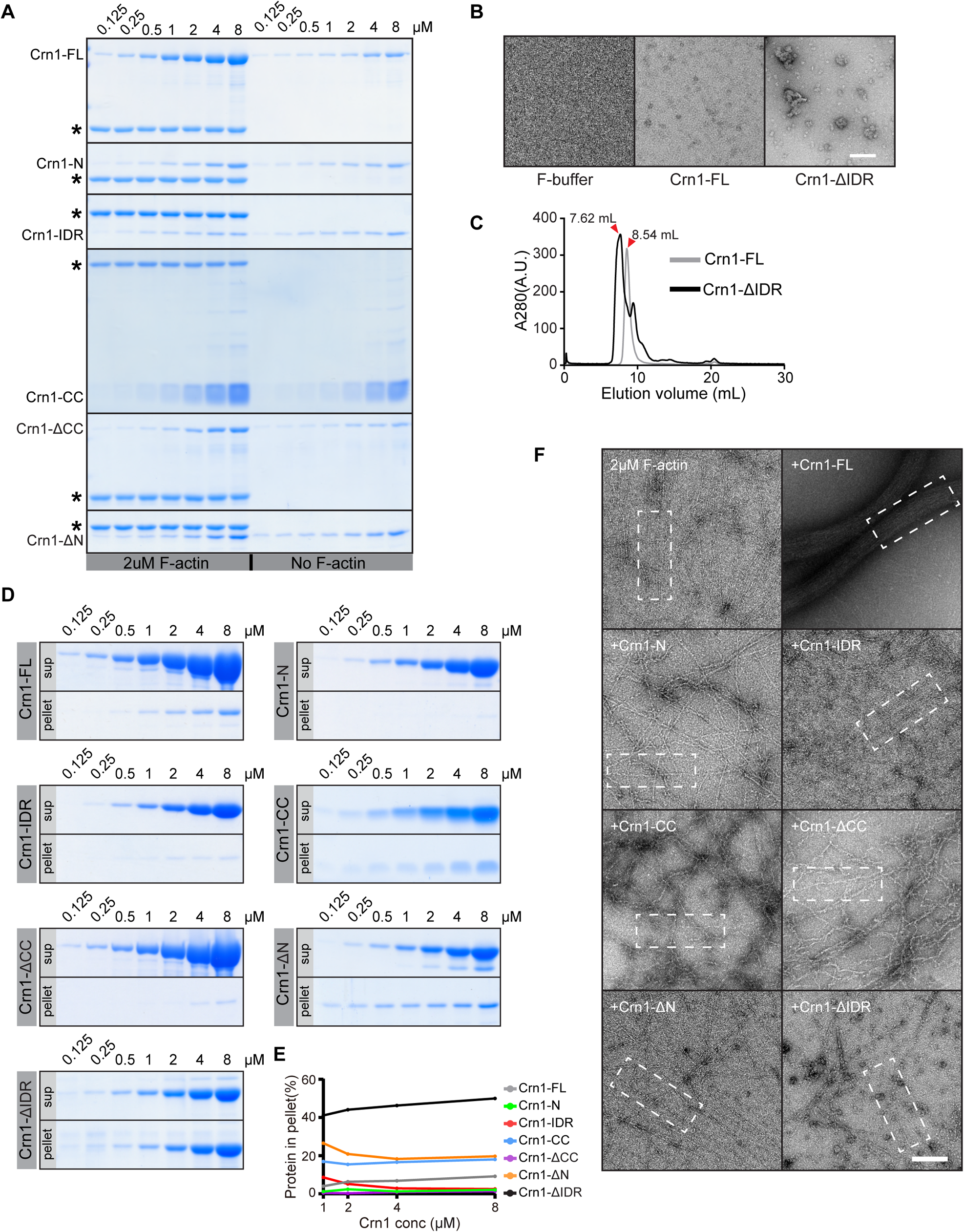
F-actin binding of recombinant Crn1 variants. A. Coomassie dye-stained SDS-PAGE gel of Crn1 variants in pellet fractions with or without F-actin after high speed actin cosedimentation assay. Actin is indicated by stars.
B. Representative transmission electron microscope (TEM) images of Crn1-ΔIDR and Crn1-FL, at a concentration of 1 µM each, in F-buffer. Scale bar, 100 nm.
C. Size exclusion chromotograpgy profiles of Crn1-ΔIDR and Crn1-FL using Superdex 200 increase 10/30 column in a pH 8 Tris-buffer with 150 mM NaCl.
D. Coomassie dye-stained SDS-PAGE gel showing pellet and supernatant fractions of Crn1 variants at a range of concentrations from 0.125 to 8 µM, after centrifugation at 100,000 x *g*.
E. Comparison of Crn1 variants in pellet fraction in (D) through band densitometry analysis.
F. Representative transmission electron microscope (TEM) images of pre-assembled F-actin (2 µM) with different Crn1 variants (1 µM) in F-buffer. Scale bar, 200 nm. White dash boxes indicate images showed in Figure 1G.

**Figure 2–figure supplement 1.**
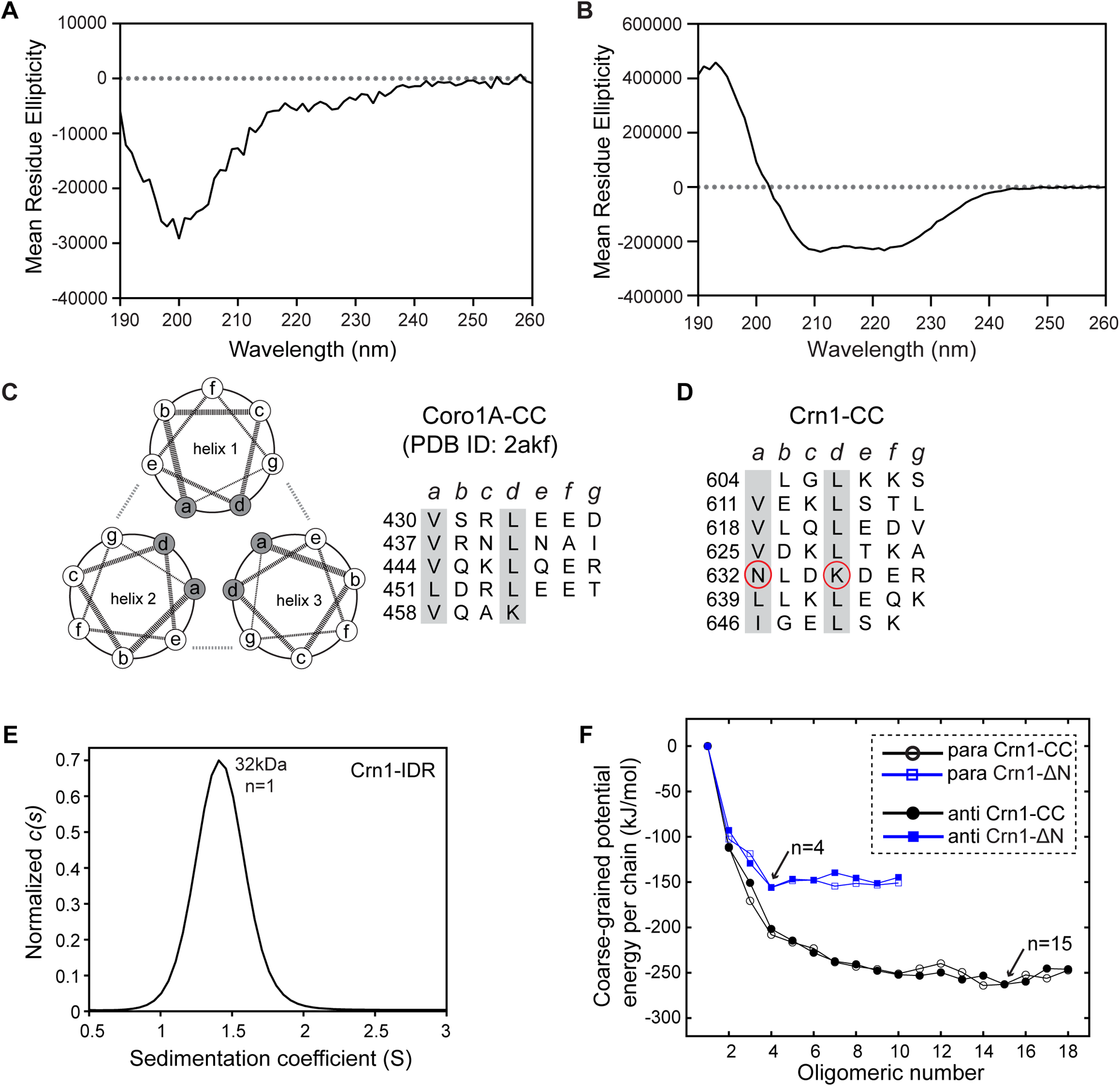
Sequence and structural analysis of Crn1-IDR and Crn1-CC. (A-B) CD spectra of Crn1-IDR (2.3 µM) (A) and Crn1-CC (3 µM) (B).
(C) Helical wheel representation (left) and sequence (right) of parallel trimeric coiled coil of murine coronin 1A C-terminus (PDB ID: 2akf). Heptad positions are labeled as a-g.
(D) Sequence of Crn1-CC assigned into heptad positions by deepcoil2, non-hydrophobic residues at the ‘a’ and ‘d’ positions are highlighted by red circles.
(E) Sedimentation coefficient distribution *c(s)* profile of Crn1-IDR (3 µM) in AUC-SV. The *c(s)* distribution was normalized to max *c(s)* in GUSSI, estimated molecular weights and oligomeric states are indicated.
(F) Potential energy per chain of Crn1-ΔN and Crn1-CC with both parallel (para) and antiparallel (anti) helical orientations varies with number of helices in coarse-grained simulations.

**Figure 3–figure supplement 1.**
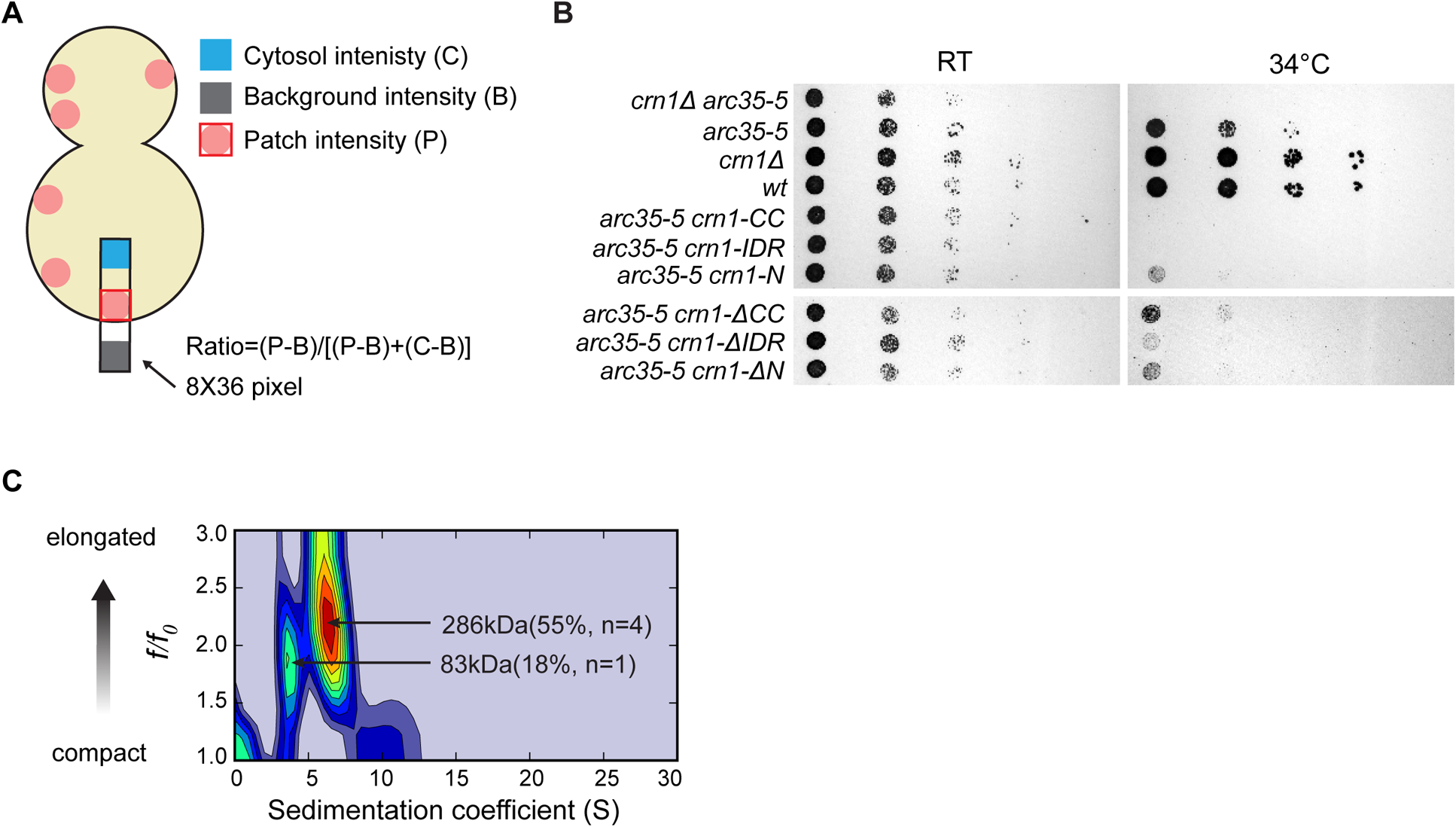
Characterization of Crn1 *in vivo* functionality. A. Cartoon diagram showing quantification of patch/total intensity ratio using the signal intensities of the cytosol (C), background (B) and patch (P) (see Methods).
B. Yeast spotting assay of Crn1 truncated variants in the genetic background of *crn1Δ arc35-5*. Cells were grown on YPD plates at the indicated temperature for 42 hours before imaging.
C. Two-dimensional size-and-shape distribution of *E.coli* purified Crn1-FL(related to Figure 2A). Different populations were determined by sedimentation coefficient (S) and frictional ratio (*f/f0*). Color in heat map indicates concentration of different populations, from lowest (blue) to highest (red). The increasing value of *f/f0* indicates elongated or non-globular species.

**Figure 4–figure supplement 1.**
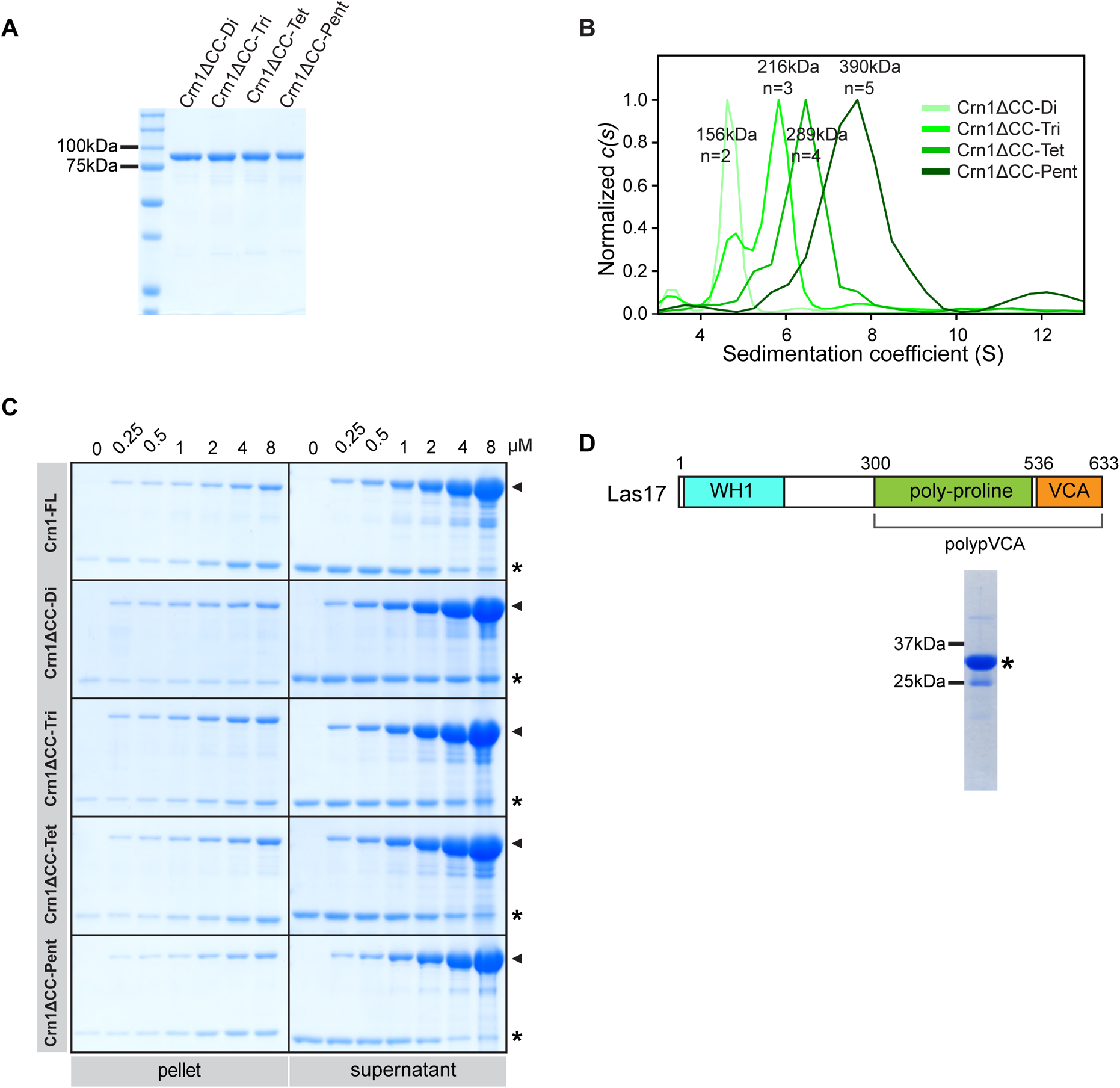
AUC verification and *in vitro* functional assays of Crn1 homo-oligomers. A. Coomassie dye-stained SDS-PAGE gel of purified recombinant Crn1 homo-oligomer variants in a similar design in Figure 3A without C-terminal tags.
B. Overlaid sedimentation coefficient distribution profiles of purified homo-oligomer Crn1 variants Crn1ΔCC-Di (7.6 µM), Crn1ΔCC-Tri (8.9 µM), Crn1ΔCC-Tet (8.7 µM) and Crn1ΔCC-Pent (9 µM) from AUC-SV experiments performed in 50 mM Tris pH 8, 150 mM NaCl. The *c(s)* distribution was normalized to max *c(s)* in GUSSI, estimated molecular weights and oligomeric states are indicated.
C. Coomassie dye-stained SDS-PAGE gel showing actin in pellet and supernatant fractions after low-speed actin cosedimentation assay in the presence of a range of concentrations of Crn1 homo-oligomer variants in buffer with physiological ionic strength of 150 mM KCl. Actin and Crn1 proteins are indicated by stars and arrowheads, respectively.
D. Domain illustration and Coomassie dye-stained gel of purified recombinant Las17-polypVCA.

**Figure 6–figure supplement 1.**
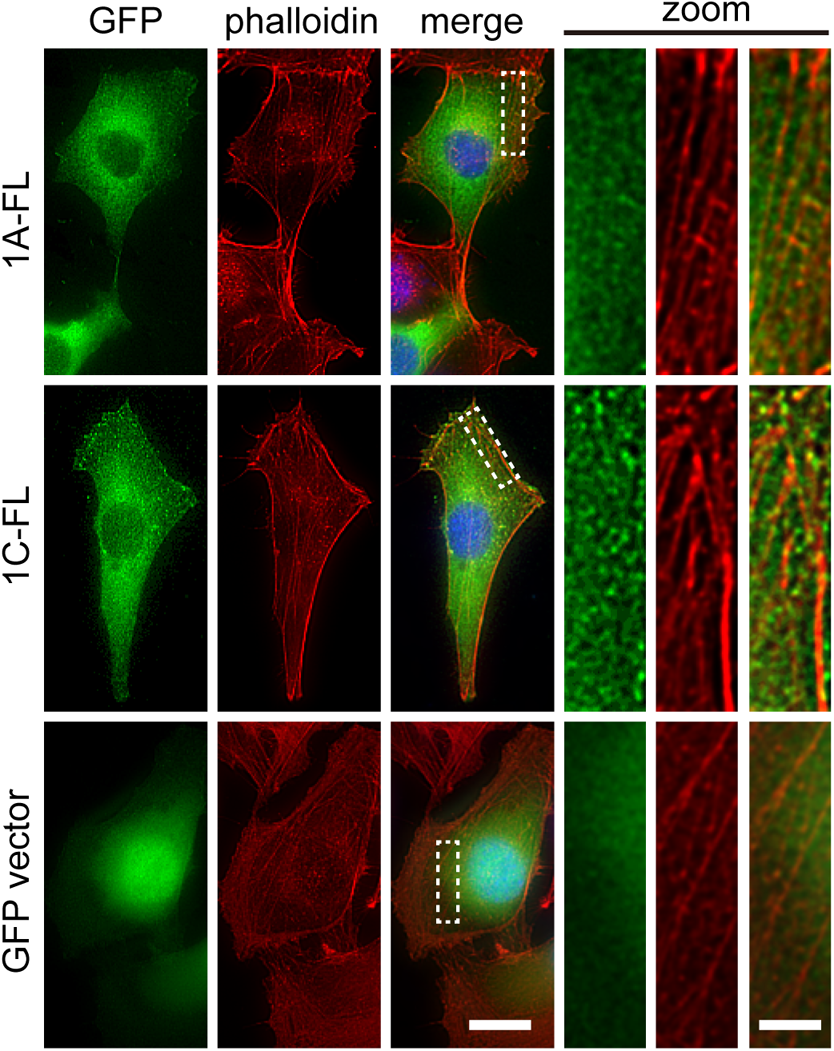
IDR roles in mammalian coronins for F-actin colocalization. Representative images of the minor population of MEF cells overexpressing full-length Coronin 1A-GFP, Coronin 1C-GFP, and empty vector control (total 1A-FL, n=78; 1C-FL, n=33; vector, n=21 cells). Zoomed images were generated from white dashed boxes. Scale bar: 20 µm (left), 5 µm (zoom).

**Figure 7–figure supplement 1.**
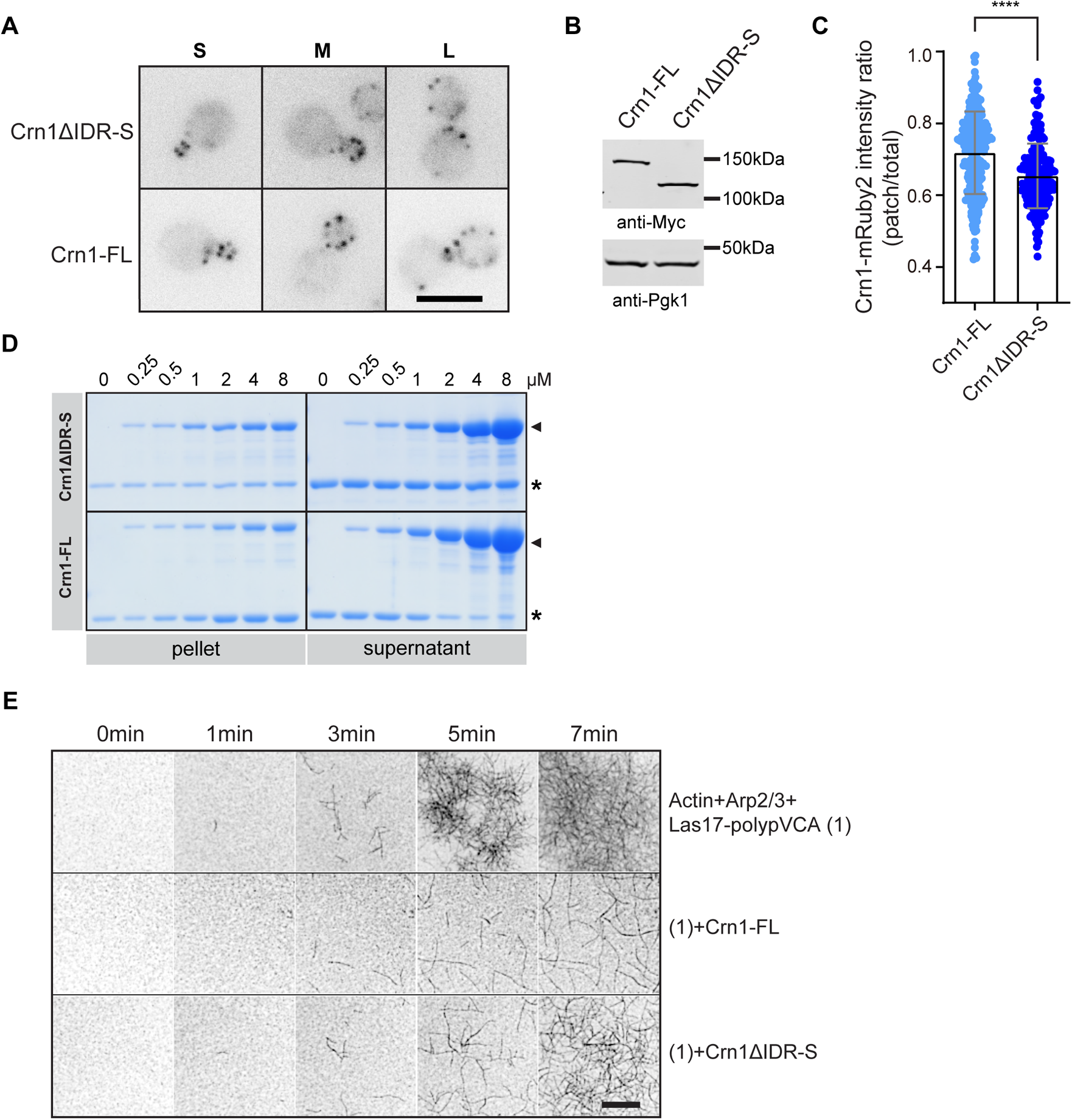
IDR tunes Crn1 function in a length-dependent manner. A. Representative fluorescent images of mRuby2-tagged Crn1-FL and Crn1ΔIDR-S. S, small-budded cell; M, middle-budded cell; L, large-budded cell. Scale bar, 5 µm.
B. Western blot detection of *in vivo* expressed mRuby2-tagged Crn1ΔIDR-S using anti-Myc antibody. The anti-Pgk1 antibody was used as a loading control.
C. Fluorescence intensity ratio(patch/total, see Methods) of mRuby2-tagged Crn1-FL (n=230 patches) and Crn1ΔIDR-S (n=189 patches) on actin patches in (A).
D. Representative Coomassie dye-stained SDS-PAGE gel of actin in the supernatant and pellet fractions after low-speed actin cosedimentation assay in the presence of a range of concentrations of Crn1ΔIDR-S or Crn1-FL proteins in buffer with physiological ionic strength of 150 mM KCl. Actin and protein are indicated by stars and arrowheads, respectively.
E. Representative time-lapse TIRF images of Arp2/3-mediated actin polymerization in the presence of indicated Crn1 variants, including 1 µM actin, 5 nM Arp2/3 complex, 25 nM Las17-polypVCA, and 200 nM Crn1 variants. Scale bar, 10 µm. Corresponding Figure 7-video 1 is available online.

**Figure 7–figure supplement 2.**
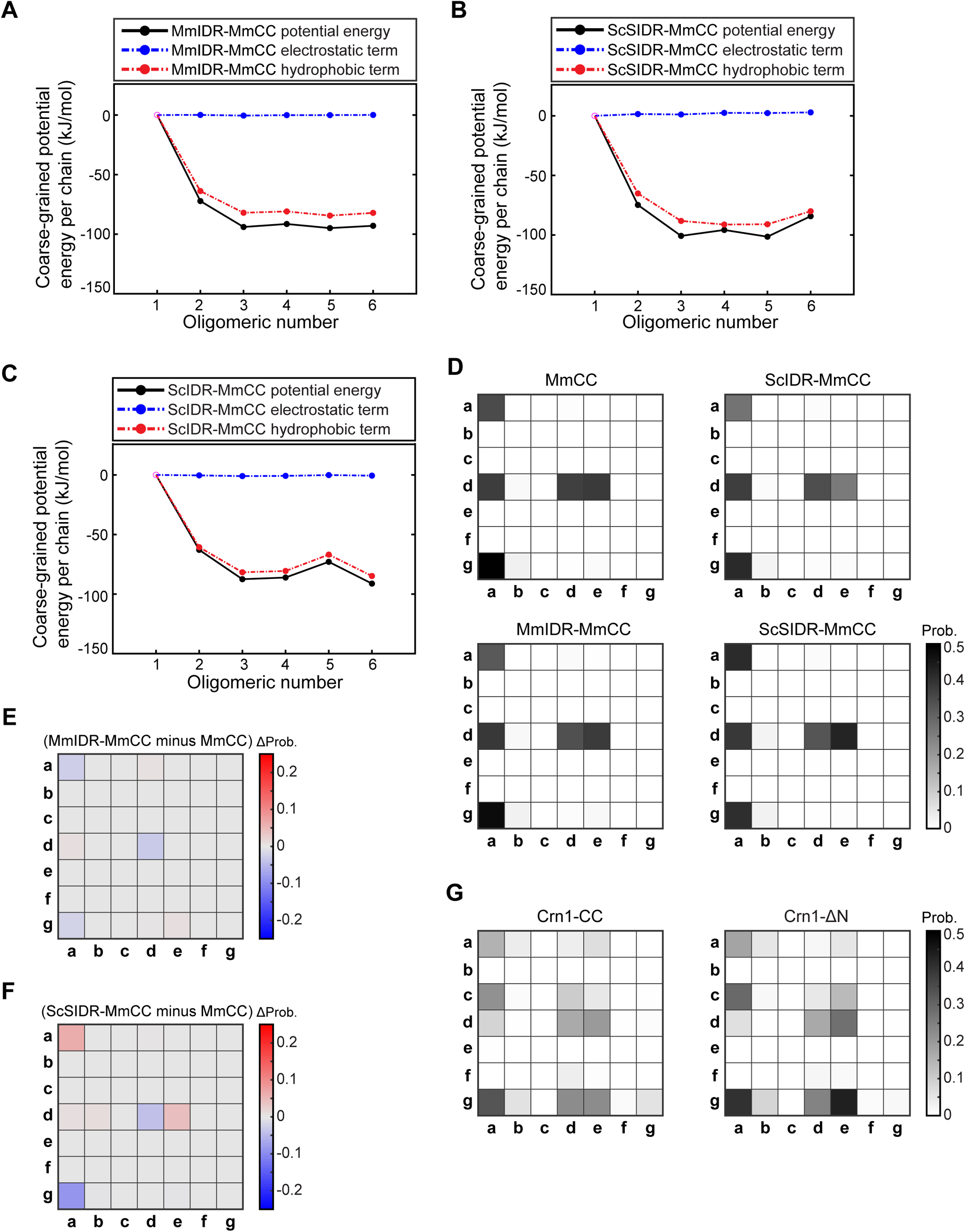
Energy landscape of coronin CCs with different IDRs from coarse-rained simulations. (A-C) Potential energy per chain of MmIDR-MmCC (A), ScSIDR-MmCC (B), and ScIDR-MmCC (C) from Figure 7H varies with number of helices in coarse grained simulations. Potential energy per chain was decomposed into hydrophobic term and electrostatic term for analysis of contribution.
(D) Motif-averaged contact maps between heptads (abcdefg) in CC region from two adjacent chains of MmCC (PDB ID:2akf), and three recombinants of MmCC and IDRs: MmIDR-MmCC, ScSIDR-MmCC, and ScIDR-MmCC, which derived Figure 7I and (E-F) below.
(E-F) Motif contact difference map between MmIDR-MmCC and MmCC (E), and between ScSIDR-MmCC and MmCC (F). Red and blue in motif contact difference map indicate increase and decrease in contact probability, respectively.
(G) Motif-averaged contact maps between heptads (abcdefg) in CC region from two adjacent chains of Crn1-CC and Crn1-ΔN, which derived Figure 7J.

**Figure 7–figure supplement 3.**
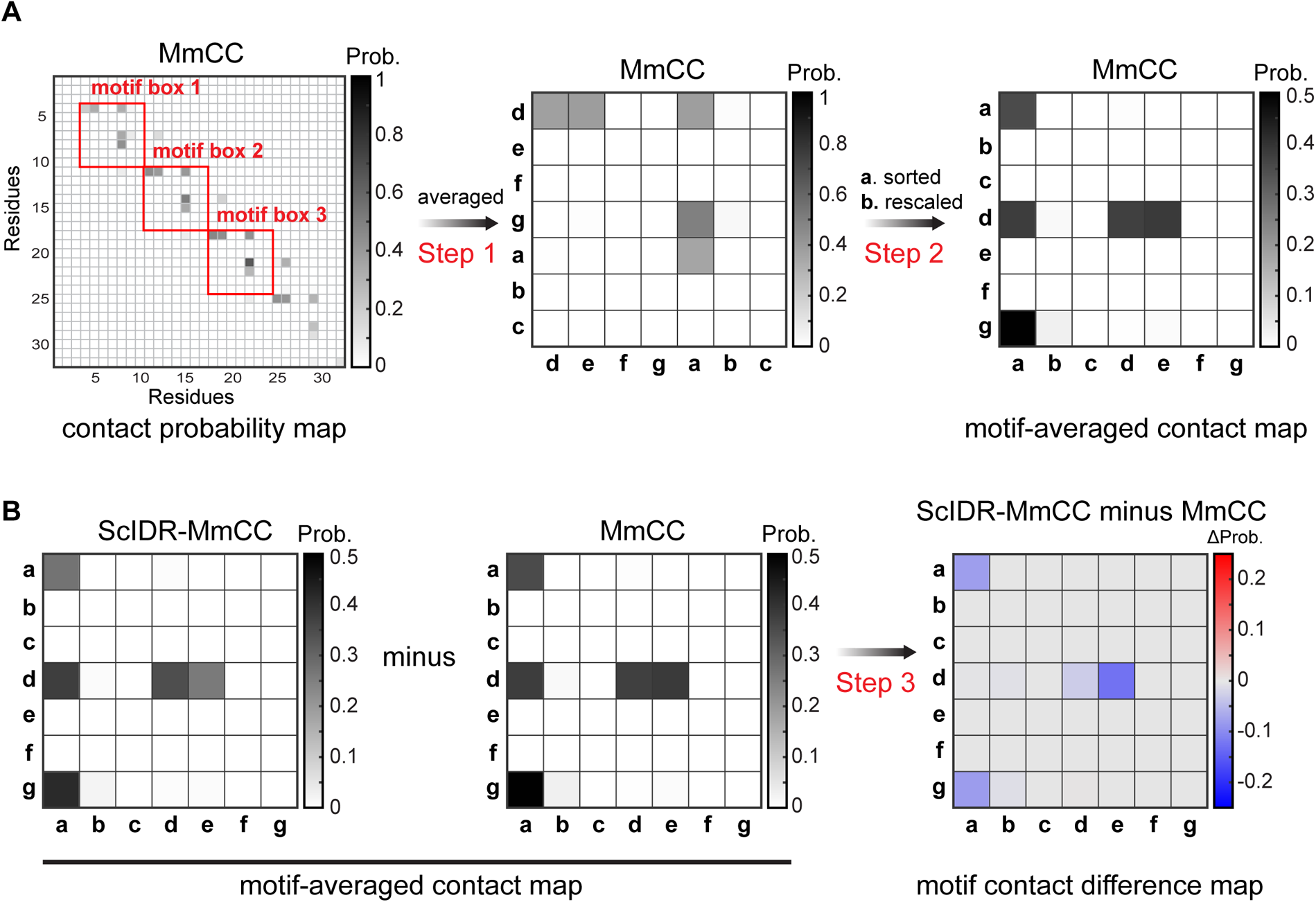
Workflow of generating motif-averaged contact map and motif contact difference maps. A. Generation of motif-averaged contact map (MmCC in this demo). The red box indicates a heptad-based motif box in a contact probability map. The motif-averaged contact map is derived from the average values of all the motif boxes in step 1 (as in the figure) followed by sorting and color rescaling in step 2.
B. Generation of motif contact difference map. With subtraction of one motif-averaged contact map (MmCC in this demo) from the other map of same type (ScIDR-MmCC in this demo), the motif contact difference map for ScIDR-MmCC minus MmCC is generated. For all the maps, the top names indicate the protein species, while the bottom ones indicate map types.

**Table S1:**
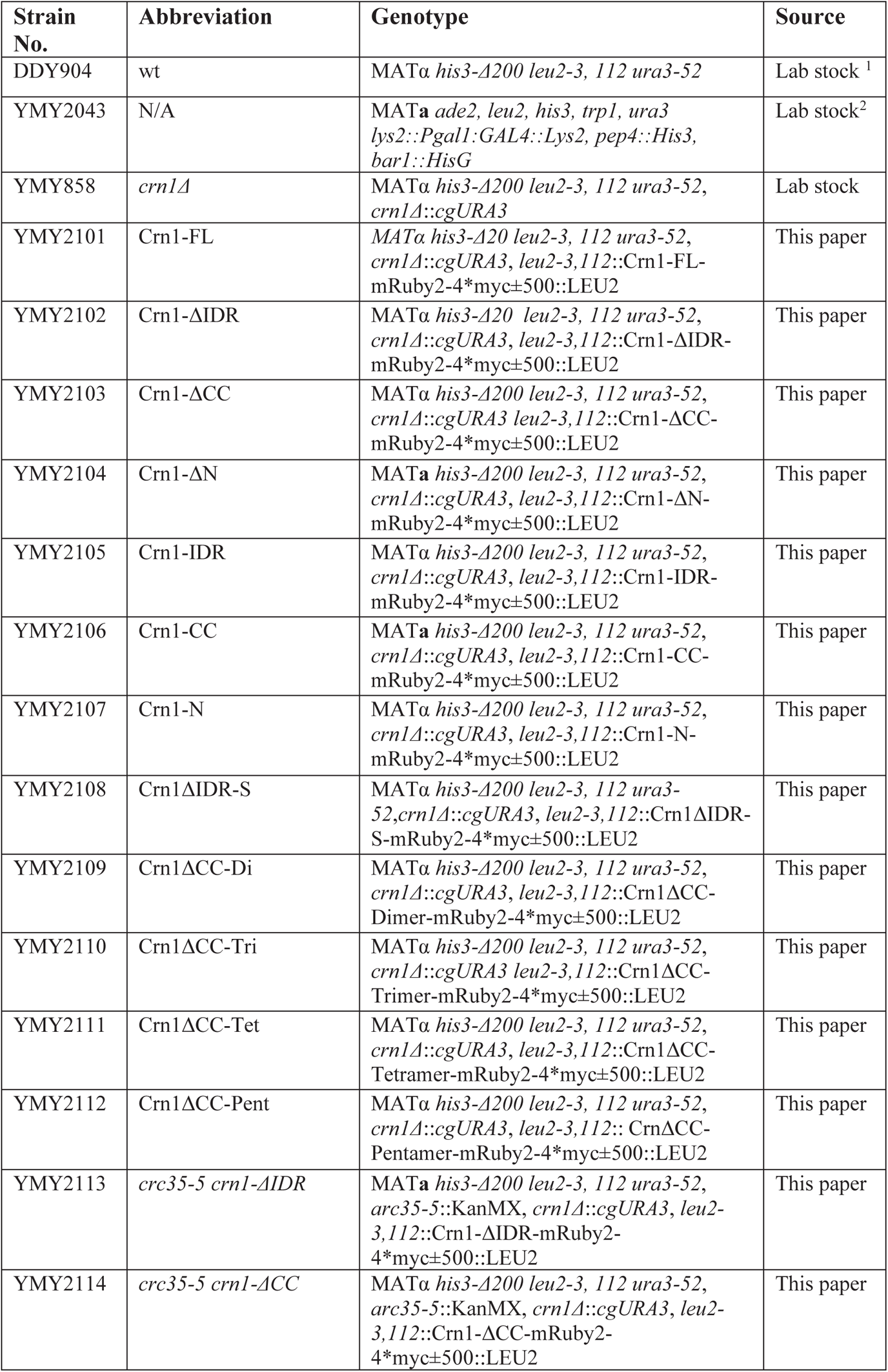
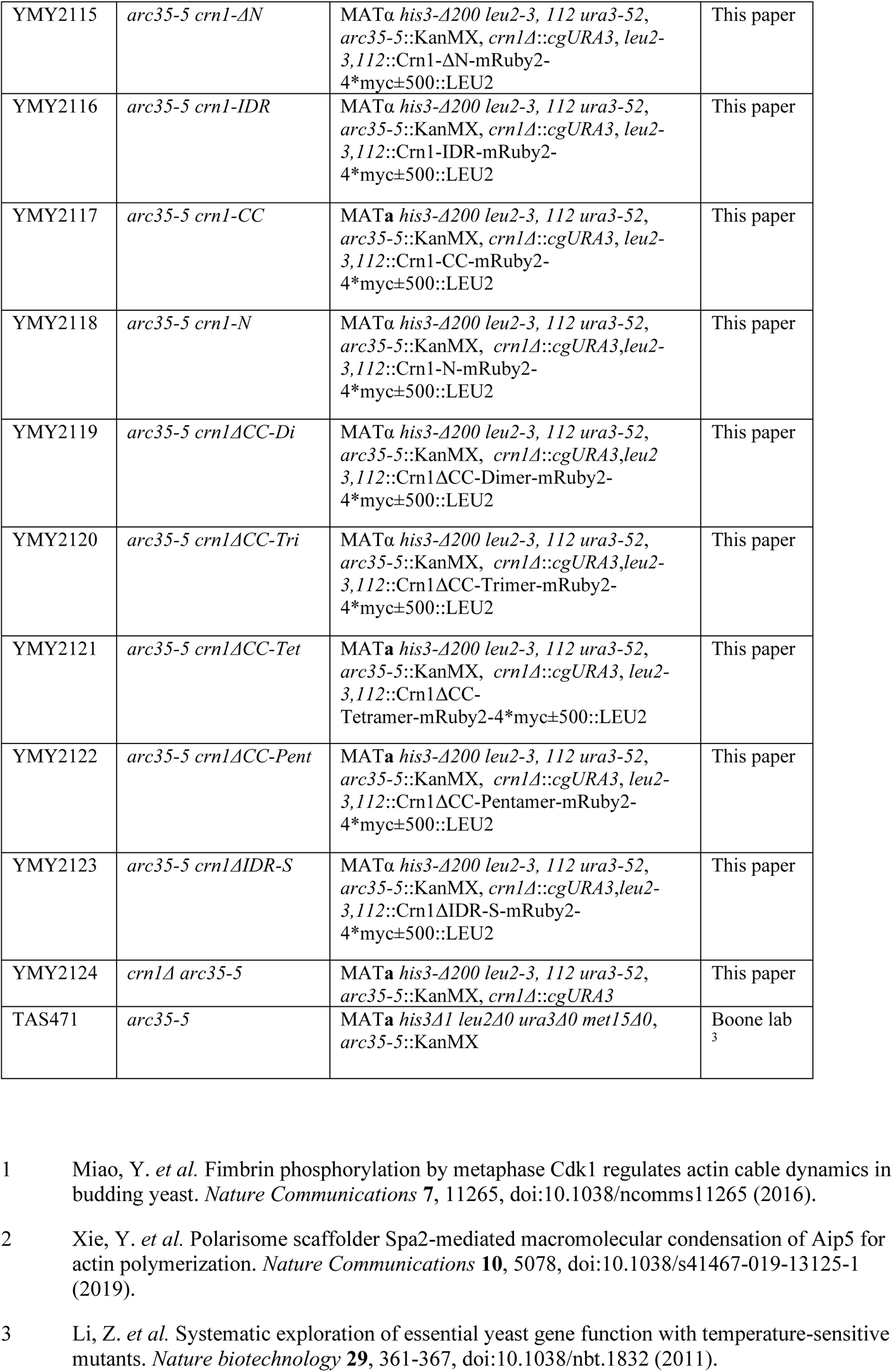
List of yeast strains used in this study.

**Table S2:** List of plasmids used in this study.

| Plasmid No. | Name | Region | Source |
| --- | --- | --- | --- |
| pMYS1501 | pRS305-Crn1-FL-mRuby2-4*myc±500 | 1-651aa of Crn1 | This paper |
| pMYS1502 | pRS305-Crn1-ΔN-mRuby2-4*myc±500 | 401-651aa of Crn1 | This paper |
| pMYS1503 | pRS305-Crn1-ΔIDR-mRuby2-4*myc±500 | 1-400+602-651aa of Crn1 | This paper |
| pMYS1504 | pRS305-Crn1-ΔCC-mRuby2-4*myc±500 | 1-604aa of Crn1 | This paper |
| pMYS1505 | pRS305-Crn1-N-mRuby2-4*myc±500 | 1-400aa of Crn1 | This paper |
| pMYS1506 | pRS305-Crn1-CC-mRuby2-4*myc±500 | 605-651aa of Crn1 | This paper |
| pMYS1507 | pRS305-Crn1-IDR-mRuby2-4*myc±500 | 401-604aa of Crn1 | This paper |
| pMYS1508 | pRS305-Crn1ΔIDR-S-mRuby2-4*myc±500 | 1-400+484-523+602-651aa of Crn1 | This paper |
| pMYS1509 | pRS305-Crn1ΔCC-Dimer-mRuby2-4*myc±500 | 1-604aa of Crn1+dimer CC sequence <sup>1</sup> | This paper |
| pMYS1510 | pRS305-Crn1ΔCC-Trimer-mRuby2-4*myc±500 | 1-604aa of Crn1+trimer CC sequence <sup>1</sup> | This paper |
| pMYS1511 | pRS305-Crn1ΔCC-Tetramer-mRuby2-4*myc±500 | 1-604aa of Crn1+tetramer CC sequence <sup>1</sup> | This paper |
| pMYS1512 | pRS305-Crn1ΔCC-Pentamer-mRuby2-4*myc±500 | 1-604aa of Crn1+pentamer CC sequence <sup>1</sup> | This paper |
| pMYS1513 | PEGFP-N1-Coro1A-FL | 1-461aa of Coro1A | This paper |
| pMYS1514 | PEGFP-N1-Coro1A-ΔIDR | 1-400+424-461 of Coro1A | This paper |
| pMYS1515 | PEGFP-N1-Coro1C-FL | 1-474aa of Coro1C | This paper |
| pMYS1516 | PEGFP-N1-Coro1C-ΔIDR | 1-400+434-474 of Coro1C | This paper |
| pMYS1517 | pSY5-Crn1-FL | 1-651aa of Crn1 | This paper |
| pMYS1518 | pSY5-Crn1-ΔN | 401-651aa of Crn1 | This paper |
| pMYS1519 | pSY5-Crn1-ΔIDR | 1-400+605-651aa of Crn1 | This paper |
| pMYS1520 | pSY5-Crn1-ΔCC | 1-604aa of Crn1 | This paper |
| pMYS1521 | pSY5-Crn1-N | 1-400aa of Crn1 | This paper |
| pMYS1522 | pSY5-Crn1-IDR | 401-604aa of Crn1 | This paper |
| pMYS1523 | pSY5-Crn1-CC | 605-651aa of Crn1 | This paper |
| pMYS1524 | pSY5-Crn1-ΔIDR-S | 1-400+484-523+602-651aa of Crn1 | This paper |
| pMYS1525 | pSY5-Crn1-ΔCC-Dimer | 1-604aa of Crn1+dimer CC sequence <sup>1</sup> | This paper |
| pMYS1526 | pSY5-Crn1-ΔCC-Trimer | 1-604aa of Crn1+trimer CC sequence <sup>1</sup> | This paper |
| pMYS1527 | pSY5-Crn1-ΔCC-Tetramer | 1-604aa of Crn1+tetramer CC sequence <sup>1</sup> | This paper |
| pMYS1528 | pSY5-Crn1-ΔCC-Pentamer | 1-604aa of Crn1+pentamer CC sequence <sup>1</sup> | This paper |
| pMYS1529 | pSY5-Las17-polypVCA | 300-633aa of Las17 | This paper |
| pMYS1530 | P634-Crn1-FL | 1-651aa of Crn1 (yeast expression) | This paper |

1 Khairil Anuar, I. N. A. *et al.* Spy&Go purification of SpyTag-proteins using pseudo-SpyCatcher to access an oligomerization toolbox. *Nature Communications* **10**, 1734, doi:10.1038/s41467-019-09678-w (2019).

**Table S3:**
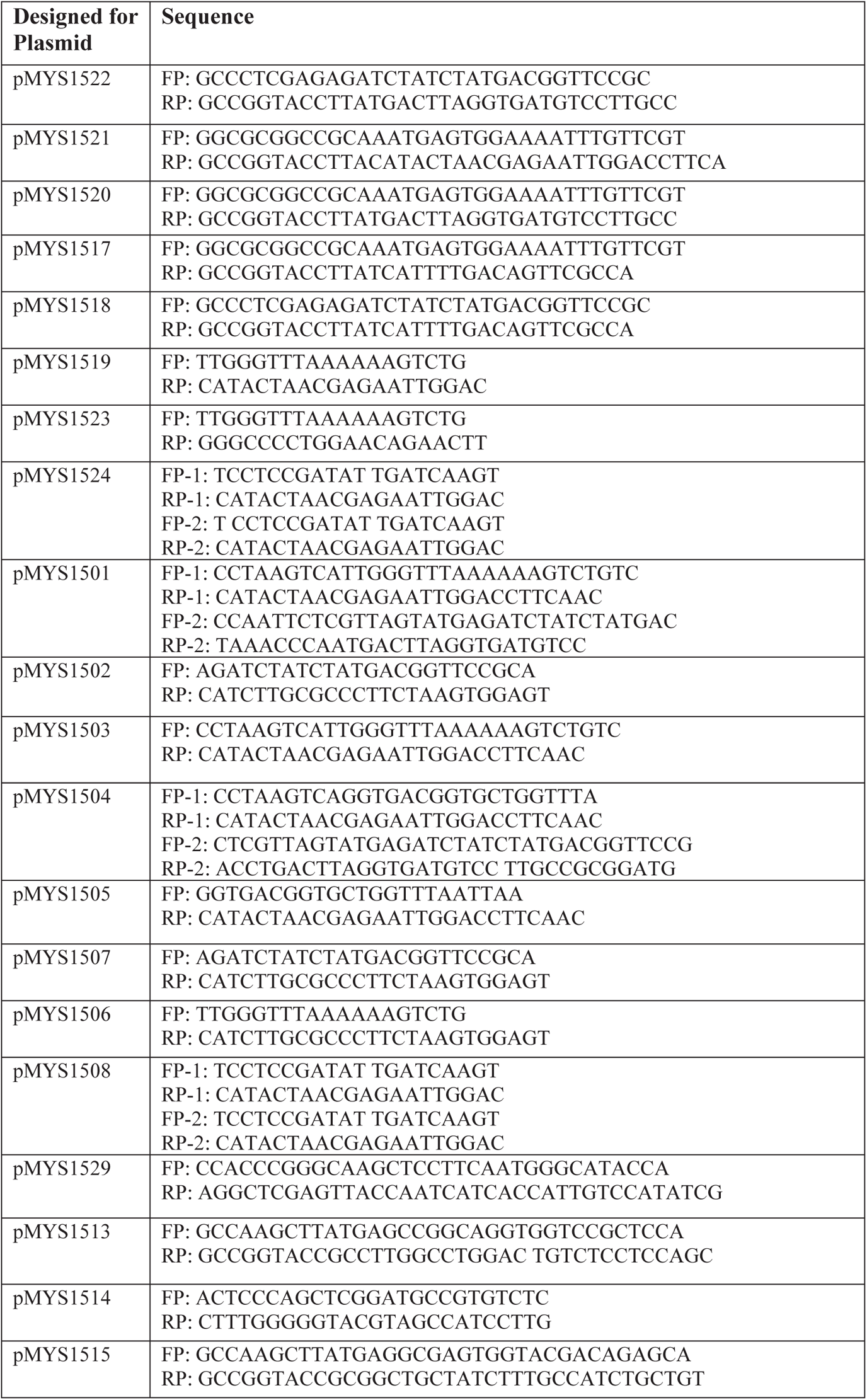
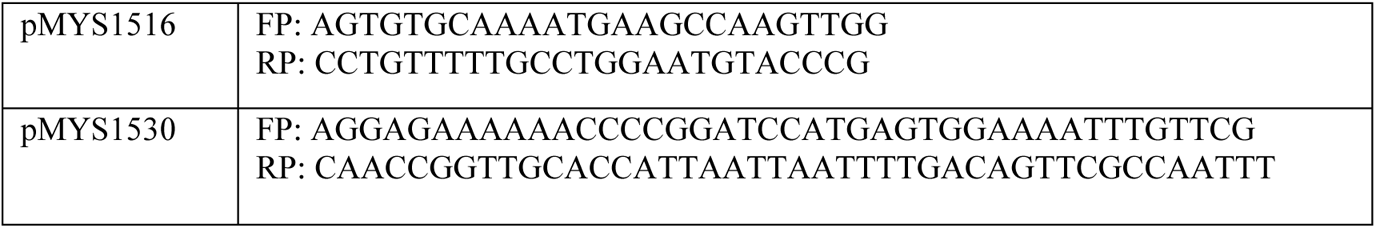
List of primers used in this paper.

## Notes

### Competing Interest Statement

The authors have declared no competing interest.

